# Lanthanide protein biosensors with a single ion-binding site

**DOI:** 10.64898/2026.08.25.747157

**Authors:** Isabella Nymann Westensee, Zhong Guo, Zhenling Cui, Chantal Ronacher, Micaela Fiorito, Alex Beliaev, Kirill Alexandrov

## Abstract

Rising demand for rare earth elements, including lanthanides (Lns), has intensified environmental pressures and supply-chain vulnerabilities, motivating the development of bio-based methods for their extraction and separation. However, the lack of high-throughput assays for analysing the selectivity of lanthanide-binding proteins remains a key bottleneck in engineering bio-based Ln-extraction systems. Here, we report the development of high-throughput assays based on Ln-responsive protein biosensors. These β-lactamase-based biosensors contain receptors with a single Ln-binding site derived from either lanmodulin or the AI-designed protein RF2. We established multiplexed colourimetric assays that quantify biosensor activity and selectivity *in vitro* and in the periplasm of *E. coli*. We further demonstrate that *E. coli* cells expressing these biosensors exhibit Ln-dependent survival in the presence of β-lactam antibiotics. These platforms enable large-scale testing of Ln biosensors and Ln-binding proteins.

## Introduction

Rare earth elements (REEs), which include the lanthanide (Ln) series, are widely used in electronics, medicine, energy storage and energy generation. Conventional physicochemical approaches to lanthanide separation are resource-intensive and environmentally harmful,^1,2^ creating substantial barriers to expanding Ln supply.^3-5^ Consequently, global Ln shortages have become a national security concern for advanced economies. The discovery of Ln-utilising methylotrophic bacteria^6^ and their selective Ln-binding proteins, lanmodulins (LanM),^7,8^ raised hopes for cleaner and more efficient Ln bioextraction technologies. This is supported by demonstrations of LanM-based separation of Lns from non-REE ions in solution,^9^ chromatographic Ln separation using LanM immobilised on magnetic nanoparticles,^10^ LanM displayed on the surface of the fungus *Yarrowia lipolytica*,^*11*^ and LanM-functionalised silk sponges.^12^ Separation of heavy and light Lns has also been demonstrated using LanM-immobilised microbeads,^13^ while neodymium/dysprosium separation was achieved using LanM from *Hansschlegelia quercus*.^14^

Despite these advances, industrial Ln-separation processes remain distant because of limitations at both the component and systems levels.^15^ It remains unclear whether protein composition and structure can be tuned to achieve the selectivity required to separate adjacent Lns. To address this question, Zhang et al. used LanM EF-hand 2 (EF2) as a template to rationally design a peptide for immobilisation on magnetic agarose beads; the resulting material selectively enriched Sc(III) in the presence of non-REE ions and Eu(III) and Tb(III).^16^ Verma *et al*. showed that varying the isoleucine residue in an EF1 loop-derived peptide altered selectivity for La(III), Ce(III), Pr(III) and Nd(III).^17^ In a non-LanM-based approach, Khoury *et al*. used phage-assisted continuous evolution of a calmodulin-derived peptide library, coupled through lanthanide-mediated protein-protein interactions, to transform a non-selective scaffold into a protein with REE-binding selectivity.^18^ More recently, Diep *et al*. developed the ‘SpyTag-Catcher Immobilization of Lanmodulin for Assaying Metal-Binding Selectivity’ (SpyCI-LAMBS) assay and measured the intra-REE selectivity of 621 LanM orthologues, revealing eight distinct selectivity clusters.^19^ These studies demonstrate that protein selectivity can be tuned, but do not establish its achievable limit.

One of the key limitations in both prospecting for and engineering Ln-binding proteins is the lack of methods for high-throughput analysis of binding selectivity. Current methods rely either on inductively coupled plasma mass spectrometry (ICP-MS) or on binding-induced luminescence from certain members of the Ln series, such as terbium. Neither approach adequately supports protein-engineering campaigns: ICP-MS is difficult to scale, whereas luminescence-based measurements are limited to a subset of Lns. Protein biosensors that couple Ln binding to a measurable catalytic output therefore offer an attractive alternative that can be assessed at high throughput. They can also serve as reporters in competitive assays to measure the Ln-binding affinity and selectivity of other molecules.

Previously, we reported a TEM-1 β-lactamase-LanM chimera (BLA-LanM) that couples Ln binding to BLA activity, which can be monitored using colourimetric or electrochemical assays or an *E. coli* survival assay.^20^ This platform is useful for monitoring Lns in complex mixtures; however, its use in engineering Ln-binding selectivity is complicated by the presence of multiple Ln-binding EF-hands and the complex cooperative interactions among them. Simultaneous evolution of multiple binding sites is beyond the reach of current protein-engineering and evolutionary methods. Although this complexity can be reduced by ablating individual EF-hands, the resulting biosensors have low dynamic ranges and are not ideal for engineering Ln specificity.^21^

Here, we report the construction of high-throughput assays based on Ln biosensors containing a single Ln-binding site. We demonstrate that β-lactamase-based protein biosensors can be used to analyse the selectivity of both natural and AI-designed Ln-binding proteins.

## Results and Discussion

### Functional Analysis of Protein Biosensors with LanM-Derived Receptors

To reduce the complexity of Ln-binding proteins and couple Ln binding to an easily quantifiable output, we used previously reported Ln-controlled LanM-BLA chimeric biosensors^20^. Given the allosteric interactions among the Ln-binding EF-hands and previous results showing that EF-hand ablation reduced the biosensor’s dynamic range, we hypothesised that circular permutation of LanM variants could improve allosteric coupling. We therefore designed five circularly permuted (CP) variants of each of three single-EF-hand LanM variants—LanMΔEF1,2,4, LanMΔEF1,3,4 and LanMΔEF2,3,4—and inserted them at position 41 of TEM-1 BLA. The resulting open reading frames were expressed in *E. coli*, together with the non-permuted variants BLA-LanMΔEF1,3,4 and BLA-LanMΔEF2,3,4. The recombinant proteins were purified, and their enzymatic activity was assessed in the presence or absence of 500 nM LaCl_3_ by monitoring hydrolysis of the chromogenic β-lactamase substrate UW154 (Supplementary Figure 1). Although most designs showed LaCl_3_-dependent activity, the most promising switches in terms of dynamic range and enzymatic activity were BLA-LanMΔEF1,2,4, BLA-LanMΔEF1,3,4, BLA-LanMΔEF2,3,4 and a CP variant of BLA-LanMΔEF1,2,4, designated BLA-cpLanMΔEF1,2,4-1 (Supplementary Figure 2 and Supplementary Table 1). The response times of these four switches to 500 nM LaCl_3_ were evaluated (Supplementary Figure 3); all reached full activation after 10 min.

To compare the switches in terms of apparent K_D_, dynamic range and maximum catalytic rate, we measured the responses of 15 nM of each switch to representative light, medium and heavy Lns (LaCl_3_, TbCl_3_ and LuCl_3_, respectively; Figure 1c–e and Supplementary Figures 4–7). All switches bound LaCl_3_, TbCl_3_ and LuCl_3_ with nanomolar apparent affinity. However, BLA-cpLanMΔEF1,2,4-1 had relatively low maximum catalytic activity (k_obs_; Supplementary Figure 4), whereas BLA-LanMΔEF2,3,4 had lower dynamic ranges (Supplementary Figure 5) than BLA-LanMΔEF1,3,4 (Supplementary Figure 6) and BLA-LanMΔEF1,2,4 (Figure 1d and Supplementary Figure 7). We next evaluated the responses of BLA-LanMΔEF1,3,4 and BLA-LanMΔEF1,2,4 to a broader panel of REEs (LaCl_3_, PrCl_3_, NdCl_3_, EuCl_3_, DyCl_3_, TbCl_3_, HoCl_3_, TmCl_3_, LuCl_3_, YCl_3_ and ScCl_3_; Supplementary Figures 6b–l and 7b–l, respectively). BLA-LanMΔEF1,3,4 displayed apparent K_D_ values of approximately 3–15 nM (Supplementary Figure 6m) and dynamic ranges of approximately 10–30-fold (Supplementary Figure 6n). BLA-LanMΔEF1,2,4 showed slightly higher dynamic ranges of 24–30-fold (Figure 1d), comparable maximum catalytic rates (Supplementary Figure 7m), and apparent K_D_ values of approximately 2–4 nM for light and medium REEs and approximately 20 nM for heavier REEs (Figure 1e). Both switches had substantially lower apparent affinity for ScCl_3_, with K_D_ values of approximately 80 nM for BLA-LanMΔEF1,3,4 (Supplementary Figure 6l,m) and approximately 300 nM for BLA-LanMΔEF1,2,4 (Supplementary Figure 7l and Supplementary Table 1). We then compared these selectivity profiles with that of our previously reported biosensor containing wild-type (WT) LanM^20^. In contrast to the single-EF-hand variants shown in Figure 1e, BLA-LanM displayed a relatively uniform apparent K_D_ profile across the tested REEs (Supplementary Figure 8). BLA-LanM also showed lower affinity for ScCl_3_ than for the other tested REEs, but its apparent K_D_ of 55 nM (Supplementary Figure 8l,m) was lower than those of BLA-LanMΔEF1,2,4 and BLA-LanMΔEF1,3,4 (Supplementary Table 1).

**Figure 1.**
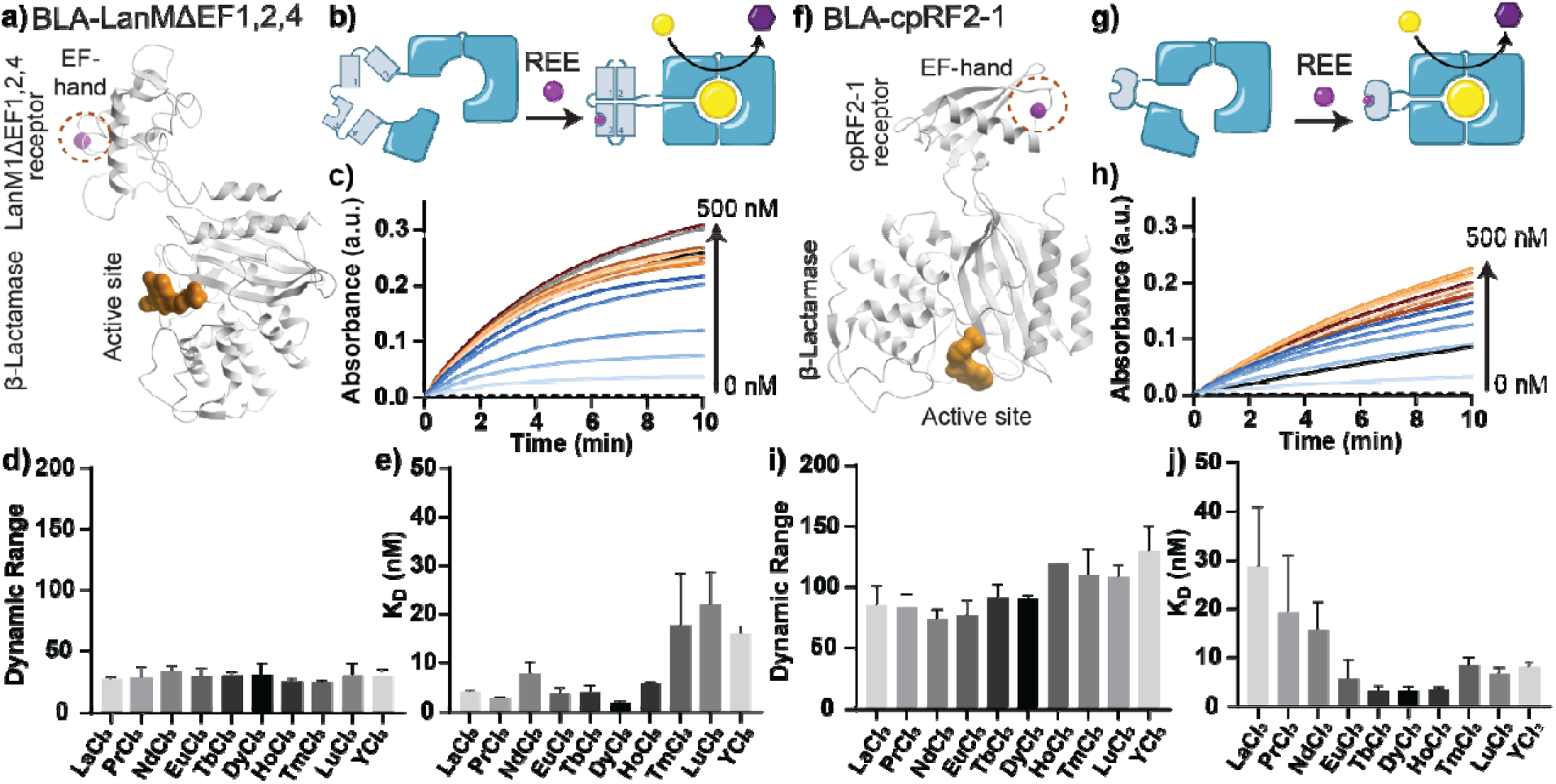
Characterisation of protein switches containing a single Ln-binding site. a) Ribbon representation of a model of the BLA-LanMΔEF1,2,4 chimera. The yttrium ion is shown as a magenta sphere, and the β-lactamase active site is marked by an inhibitor imported from PDB entry 6C79 and shown as a gold molecular surface. b) Schematic illustrating how ligand binding to BLA-LanMΔEF1,2,4 converts it from inactive to an active state, resulting in the production of hydrolysed UW154, which absorbs at 520 nm. c) Representative absorbance traces recorded after 15 nM solution of BLA-LanMΔEF1,2,4 was incubated for 10 min with different concentrations of LaCl_3_ and mixed with 50 µM UW154. d) Dynamic ranges determined from titrations such as that shown in c) for the indicated REEs. e) Apparent K_D_ values determined from the same titration data. f) Ribbon representation of a model of the BLA-cpRF2-1 chimera, displayed as in a). g) Schematic of the proposed response mechanism of BLA-cpRF2-1 upon Ln binding. h) Representative absorbance traces recorded after 10 nM BLA-cpRF2-1 was incubated for 20 min with the indicated concentrations of TbCl_3_ and mixed with 50 µM UW154. i) Dynamic ranges and j) apparent K_D_ values determined for BLA-cpRF2-1 using titrations such as that shown in h).

Taken together, these results identify BLA-LanMΔEF1,2,4 as the most promising candidate for developing a biosensor-based, high-throughput assay of Ln selectivity. It provides a favourable dynamic range of approximately 30-fold while maintaining catalytic rates comparable to those of the other switch variants. Unlike the WT LanM-based biosensor, which has comparable affinity for most members of the Ln series, BLA-LanMΔEF1,2,4 preferentially binds light over heavy Lns. It therefore provides a potential starting point for engineering REE selectivity.

### Functional Analysis of β-Lactamase-Based Chimeric Switches with EF-Hand-Based Miniprotein Receptors

Although we constructed a functional biosensor containing a single lanthanide-binding site, its receptor was structurally redundant because it retained non-functional EF-hands that were expected to remain disordered even in the presence of ligand. We therefore sought to create a minimal Ln receptor containing only one binding site. We used the recently reported RF1 and RF2 miniproteins, which were generated by hallucinating new structures around LanM EF-hand 2.^22^ The RF1 and RF2 sequences were inserted at position 197 of BLA to generate BLA-RF1 and BLA-RF2. Five CP variants of each receptor were also designed and inserted into BLA. The constructs were recombinantly expressed and purified, and their performance was evaluated (Supplementary Figures 9 and 10). BLA-RF1 was the best-performing RF1-derived switch, whereas all but one of its CP variants lacked catalytic activity (Supplementary Figure 10a). The opposite trend was observed for RF2-derived constructs: BLA-RF2 showed no lanthanide-dependent activity, whereas two CP variants, BLA-cpRF2-1 and BLA-cpRF2-5, displayed lanthanide-dependent activity with lower background activity than BLA-RF1 (Supplementary Figure 10b). We selected BLA-cpRF2-1 for further characterisation because it had higher catalytic activity than BLA-cpRF2-5 (Supplementary Figure 10b).

BLA-cpRF2-1 reached maximum catalytic activity after a 20-min incubation with LaCl_3_ (Supplementary Figure 11). We then titrated 10 nM BLA-cpRF2-1 with increasing concentrations of different Lns. The biosensor displayed a dynamic range of up to 120-fold and apparent K_D_ values of 5–30 nM (Figure 1h–j, Supplementary Figures 12b–m and Supplementary Table 1). Interestingly, BLA-cpRF2-1 displayed an Ln-selectivity profile opposite to that of BLA-LanMΔEF1,2,4 (Figure 1e,j) and LanM proteins generally^19^. BLA-cpRF2-1 showed only minimal activation by ScCl_3_, which could not be quantified reliably (Supplementary Figure 12l). Co-incubation of BLA-cpRF2-1 with TbCl_3_ and ScCl_3_ confirmed that this behaviour did not result from switch inhibition, but instead reflected a genuine lack of activation by ScCl_3_ (Supplementary Figure 13). These results indicate that the structure and context of an EF-hand substantially influence its selectivity and affinity for Lns.

We next measured the enzymatic activity of BLA-LanMΔEF1,2,4 and BLA-cpRF2-1 after incubation with micromolar concentrations of LaCl_3_, TbCl_3_ or LuCl_3_ (Supplementary Figure 14). BLA-LanMΔEF1,2,4 activity decreased above 1 µM LaCl_3_ or 500 nM TbCl_3_ or LuCl_3_, whereas BLA-cpRF2-1 activity decreased above 500 nM LaCl_3_ or 100 nM TbCl_3_ or LuCl_3_. We previously observed similar biphasic behaviour for BLA-LanM and BLA-LanEF1,2,3,4, suggesting that Ln-mediated inhibition is largely independent of the Ln-binding receptor^20^. We also assessed responses to other metal ions. As expected, neither BLA-LanMΔEF1,2,4 nor BLA-cpRF2-1 was activated by Ca^2+^, Mn^2+^, Fe^3+^ or Al^3+^ at concentrations up to 2.5 µM (Supplementary Figure 15). BLA-LanMΔEF1,2,4 responded to Cu^2+^ with an apparent K_D_ of 10 ± 3 nM and a modest dynamic range of approximately 3 (Supplementary Figure 15a(vi)), whereas BLA-cpRF2-1 showed no response to Cu^2+^ (Supplementary Figure 15b(v)). We assessed the thermostability of both switches by incubating them at 4, 18, 37 or 60 °C for up to 300 min before a 10 min incubation with LaCl_3_ or TbCl_3_ and addition of the chromogenic substrate. Both proteins retained catalytic activity after incubation at 4– 37 °C for up to 300 min; however, BLA-LanMΔEF1,2,4 showed progressively higher background activity with increasing preincubation time (Supplementary Figure 16a). By contrast, BLA-cpRF2-1 maintained low background activity even after overnight incubation at 4 °C (Supplementary Figure 16b).

### Development of an *In Vivo* Assay for Analysing Ln-Controlled Protein Switches

To establish a high-throughput screening method for Ln-controlled protein switches, we tested whether the activity of BLA-LanMΔEF1,2,4 and BLA-cpRF2-1 could be monitored directly in *E. coli* cells. We constructed *E. coli* strains that inducibly produced each switch in the periplasm. After 20 h of expression, the cells were pelleted, and BLA activity was evaluated in both the supernatant and the resuspended cell pellet using a 96-well-plate format (Figure 2a and Supplementary Figures 17a and 18a). Activity in the supernatant was assessed either by monitoring the change in absorbance of the β-lactamase substrate UW154 with a plate reader (Supplementary Figures 17b(i) and 18b(i)) or by imaging the 96-well plate with a ChemiDoc system (Supplementary Figures 17b(ii) and 18b(ii)).

**Figure 2.**
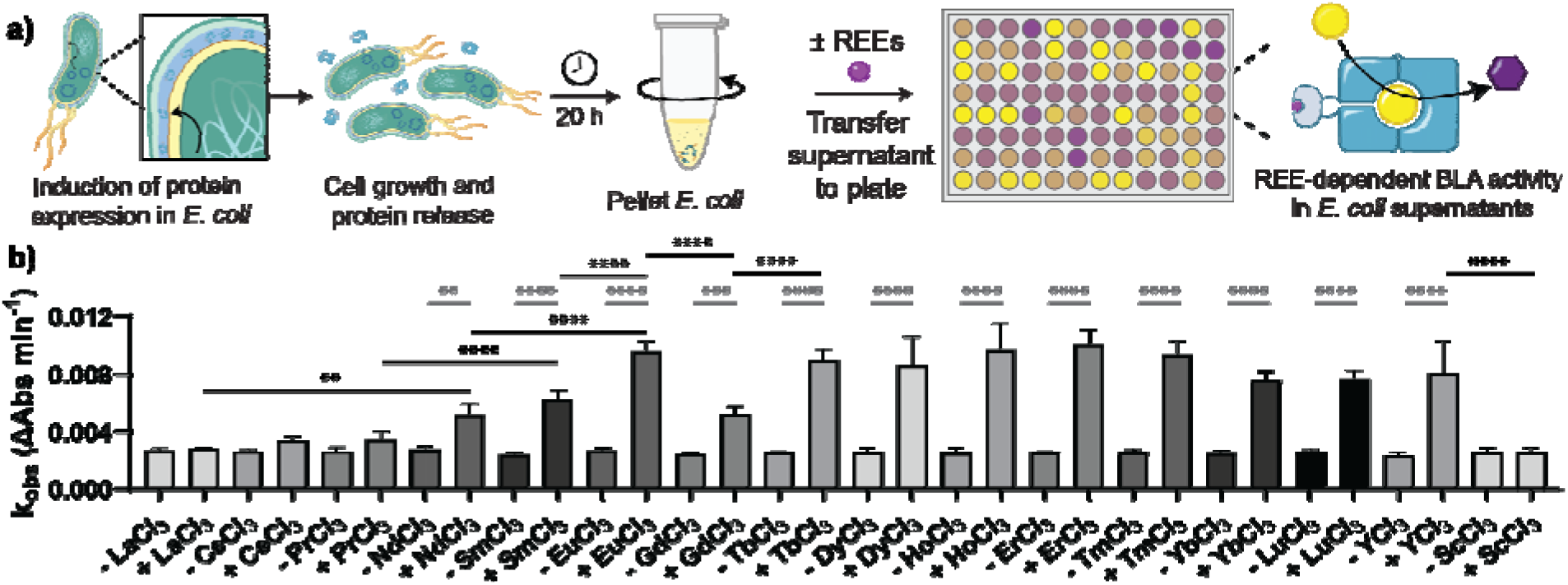
Analysis of protein-switch activity in *E. coli* expressing BLA-cpRF2-1. a) Schematic showing induction of periplasmic BLA-cpRF2-1 expression in *E. coli*, followed by cell growth, release of some protein into the medium and retention of the remainder within the cells. After 20 h, the cells were pelleted and the enzymatic activity of the chimeric BLA switch was analysed. b) k_obs_ values obtained by fitting the linear region of absorbance-time curves (λ_abs_ = 520 nm) after the supernatant was incubated with 500 nM of the indicated REE for 1 h and then mixed with 50 µM UW154. Values are shown for all tested REEs except PmCl_3_. *p < 0.05, **p < 0.005, ***p < 0.0005 and ****p < 0.00005; the absence of stars indicates no significant difference.

Supernatants from *E. coli* expressing BLA-LanMΔEF1,2,4 or BLA-cpRF2-1 displayed Ln-dependent BLA activity. The intra-REE activity profile of BLA-LanMΔEF1,2,4 closely reflected that measured with the purified protein (Figure 1e and Supplementary Figure 17b(iii)). The BLA-cpRF2-1 profile showed the same trend (Figure 1j and Figure 2b). No switching was observed when the supernatants were incubated with REEs for which the corresponding purified proteins display low affinity. BLA activity in pelleted *E. coli* cells could not be quantified with a plate reader because of the low signal-to-noise ratio for both constructs. However, after REE-exposed cells had been incubated with UW154 for 3 h, responses to different REEs could be detected by plate imaging the (Supplementary Figures 17c and 18c).

#### Ln-Controlled E. coli Survival Assays

BLA-based protein switches controlled by small molecules have been used extensively in bacterial survival-based antibiotic-selection assays^23,24^. As expected, *E. coli* cells expressing BLA-cpRF2-1 showed TbCl_3_-dependent growth in autoinduction medium containing 18–22 µg mL^-1^ ampicillin (Figure 3a,b). Cell densities were higher in the presence of medium and heavy lanthanides (Figure 3c). However, the light lanthanide NdCl_3_ supported growth to the same extent as EuCl_3_, TbCl_3_, HoCl_3_ and LuCl_3_. On solid autoinduction medium containing ampicillin, *E. coli* expressing BLA-cpRF2-1 grew only when micromolar concentrations of LaCl_3,_ NdCl_3_, TbCl_3_, HoCl_3_ or LuCl_3_ had been introduced into the agar before plating (Figure 3d and Supplementary Figure 19). Both liquid- and solid-medium assays therefore showed Ln-dependent growth, although the Ln-selectivity profile was more clearly resolved on solid medium (Figure 3c,d and Supplementary Figure 19).

**Figure 3.**
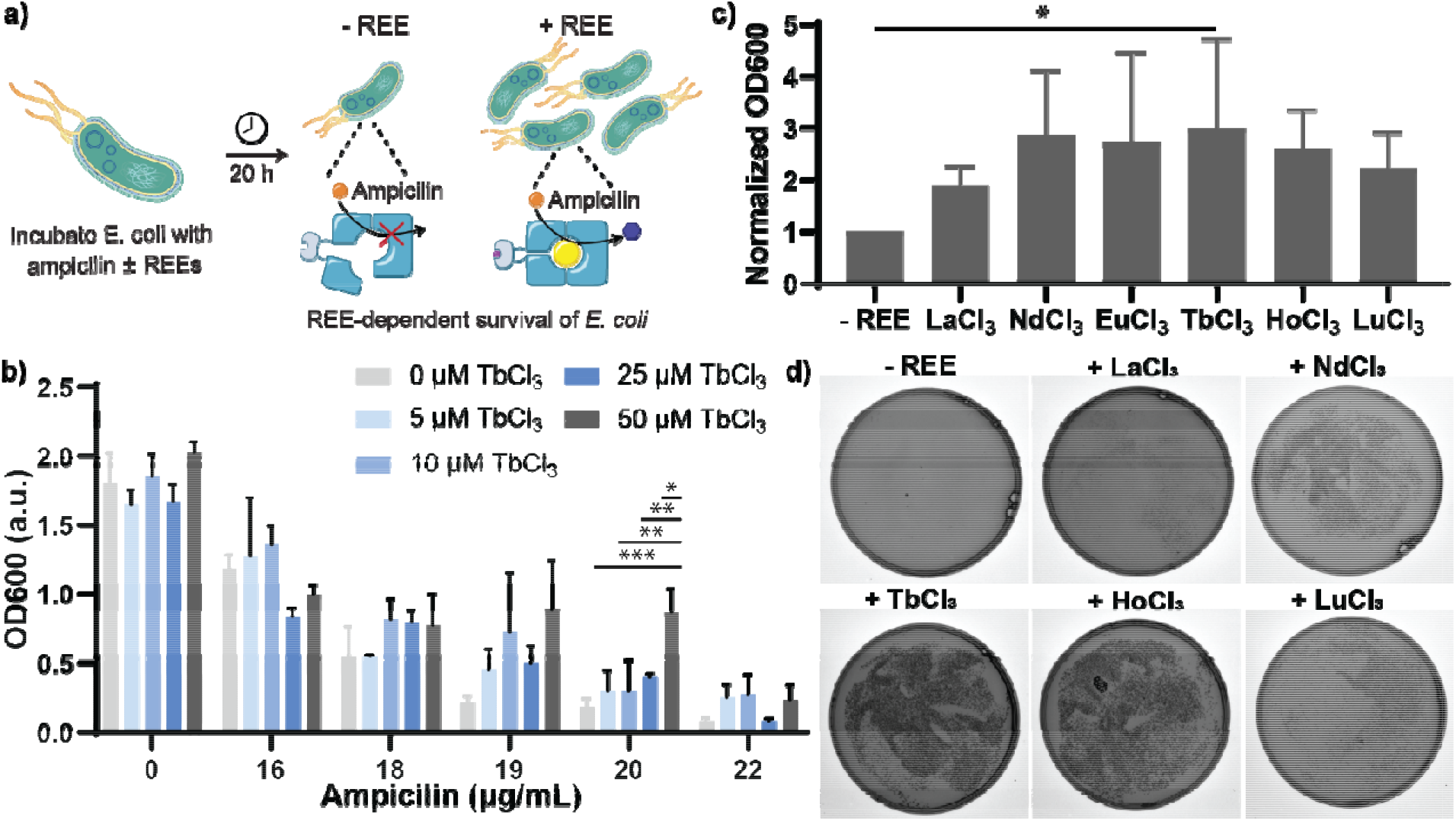
Ln-dependent survival of *E. coli* expressing BLA-cpRF2-1 under ampicillin selection. a) Schematic illustrating how formation of the Ln-bound BLA-cpRF2-1 complex promotes ampicillin cleavage and thereby increases cell survival. b) OD_600_ of *E. coli* cultures after incubation for 20 h at 30 °C with 50 µg mL^-1^ kanamycin, 0–22 µg mL^-1^ ampicillin and 0–50 µM TbCl_3_. c) OD_600_ of cultures after incubation for 20 h at 30 °C with 20 µg mL^-1^ ampicillin and 50 µM LaCl_3_, NdCl_3_, EuCl_3_, TbCl_3_, HoCl_3_ or LuCl_3_, normalised to cultures grown without REEs. d) Autoinduction agar plates containing 50 µg mL^-1^ kanamycin, 50 µg mL^-1^ ampicillin and, where indicated, 100 µM LnCl_3_. The plates were seeded with *E. coli* expressing BLA-cpRF2-1.

## Conclusions

The maximum selectivity achievable by proteins for Lns remains unknown. Ln-utilising microorganisms use Lns interchangeably and may not have experienced sufficient evolutionary pressure to discriminate between adjacent or near-adjacent elements. Protein design and directed evolution, alone or in combination, could in principle be used to test this limit; however, direct Ln-protein interaction assays such as ICP-MS are inherently low-throughput. The challenge is compounded by the multiple Ln-binding sites in natural Ln-binding proteins, which interact cooperatively and whose concurrent evolution is beyond current technical capabilities.

We therefore developed Ln biosensors that couple Ln binding at a single site to an easily quantifiable activity. We used two approaches to generate single-site receptors: ablation of selected EF-hands in LanM and use of an AI-designed Ln-binding protein^22^. Both approaches yielded functional sensors, but the latter produced the more robust design. Using these switches, we established a high-throughput platform for analysing Ln interactions with BLA-based protein switches controlled through a single REE-binding site. Importantly, switches containing a single Ln-binding domain controlled the survival of *E. coli* under antibiotic selection, opening a path towards directed evolution of Ln-binding sites. We also demonstrated that Ln selectivity and affinity can be assessed directly in living cells, providing a rapid, multiplexed method for analysing the outputs of selection campaigns. These systems can support future bioengineering applications.

## Acknowledgements

This work was supported by the Independent Research Fund Denmark (Danmarks Frie Forskningsfond) (DFF-International Postdoc grant ID: 10.46540/4257-00012) to I.N.W. This work was also supported by the DOE Office of Science through the Genomic Science Program of the Biological and Environmental Research Program under FWP 86413. Pacific Northwest National Laboratory is operated by Battelle for the U.S. Department of Energy (DOE) under Contract DE-AC05-76RL01830 to K.A. and A.B. This work was partially funded by the ARC Centre of Excellence in Synthetic Biology (CE200100029), ARC Linkage Grant LP200200916 and NHMRC Investigator Grant APP2033951, awarded to K.A. K.A. gratefully acknowledges financial support from the CSIRO-QUT Synthetic Biology Alliance and the CSIRO Synthetic Biology and Advanced Engineering Biology Future Science Platforms. This work was also supported by CAB Future Leaders seed funding and a CoESB EMCR Seed Funding Award to Z.C.

