## supplementary information for "Lanthanide protein biosensors with a single ion-binding site"

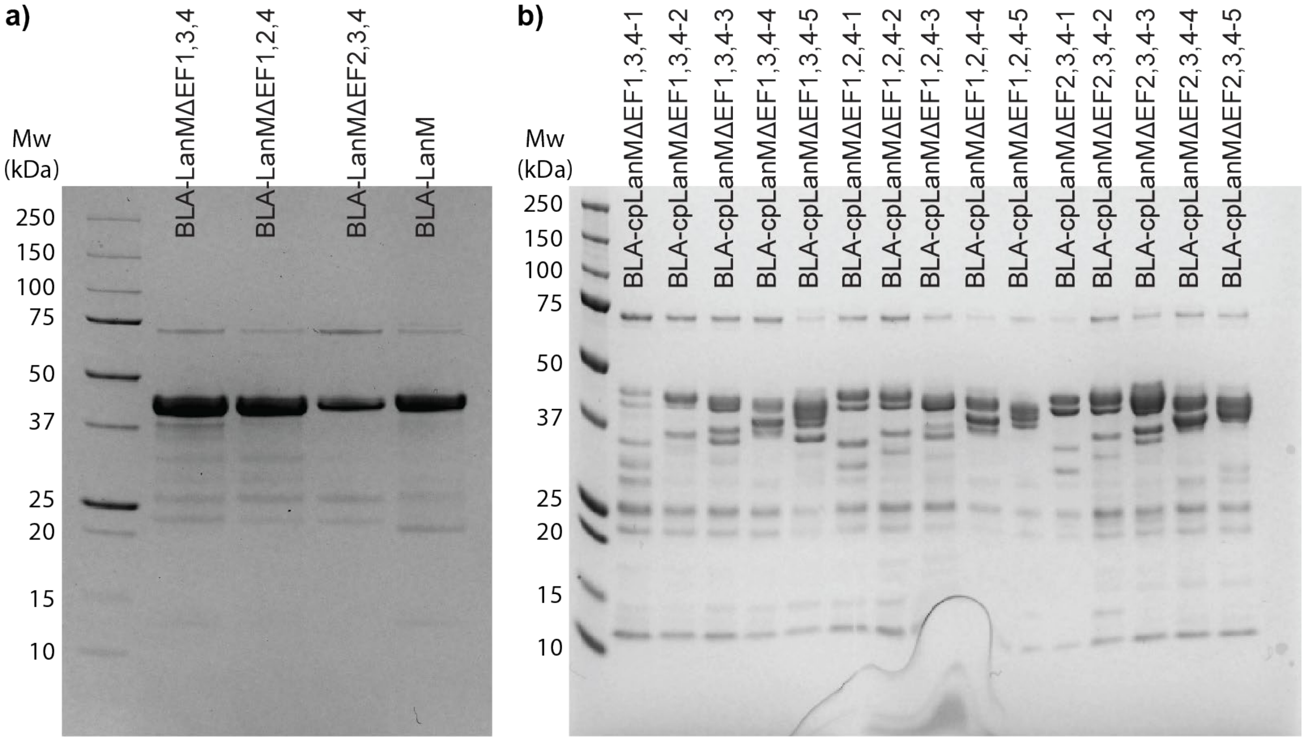


**Supplementary Figure 1**. SDS-PAGE analysis of a) BLA-LanMΔEF1,3,4, BLA-LanMΔEF1,2,4, BLA-LanMΔEF2,3,4 and BLA-LanM; and b) circularly permuted variants of BLA-LanMΔEF1,3,4 (lanes 1–5), BLA-LanMΔEF1,2,4 (lanes 6–10) and BLA-LanMΔEF2,3,4 (lanes 11–15). Each lane contained 2 µg of protein separated on a 4–12% SDS-PAGE gel and stained with Coomassie Blue.

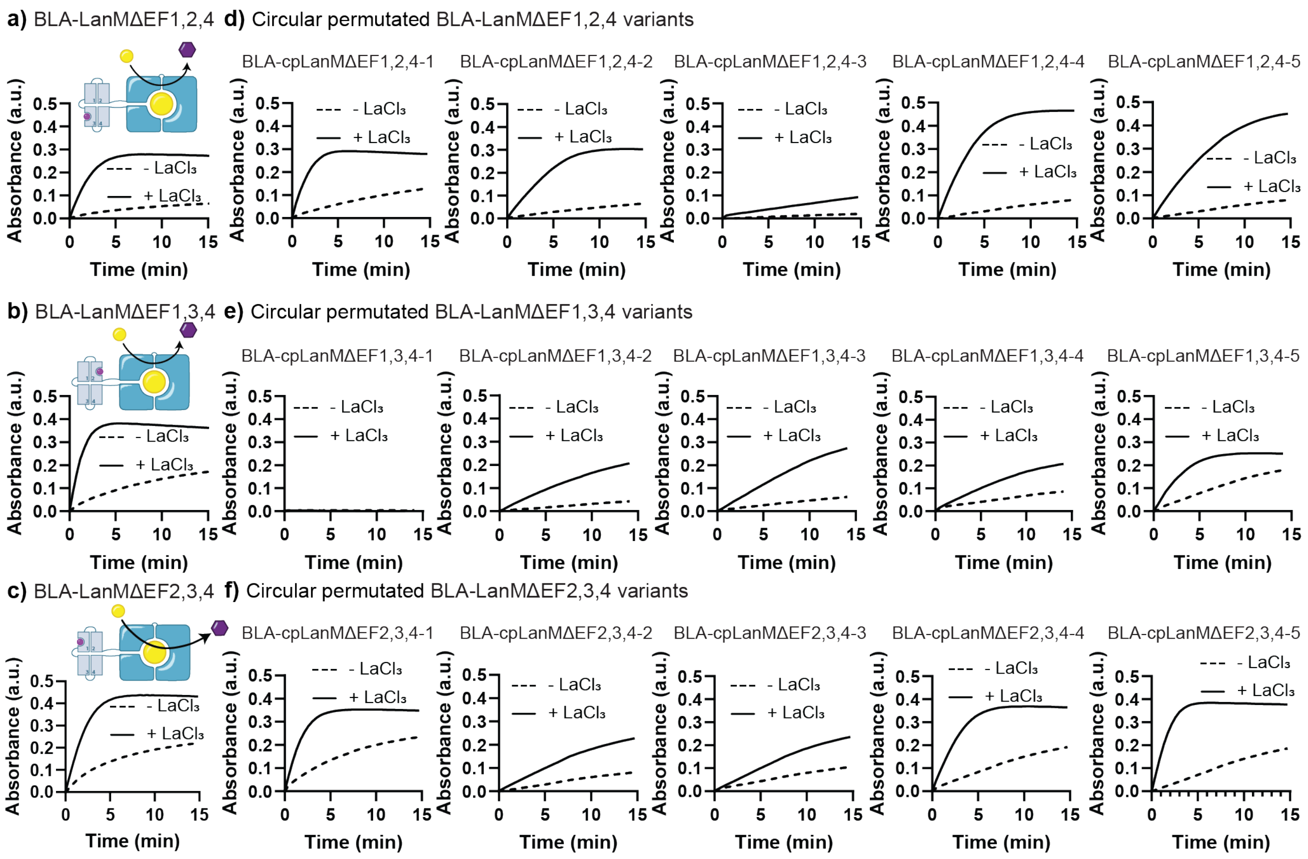


**Supplementary Figure 2.** Activity screening of a) BLA-LanMΔEF1,2,4, b) BLA-LanMΔEF1,3,4 and c) BLA-LanMΔEF2,3,4, together with the corresponding circularly permuted variants: d) BLA-LanMΔEF1,2,4, e) BLA-LanMΔEF1,3,4 and f) BLA-LanMΔEF2,3,4. Purified protein (25 nM) was incubated for 10 min with or without 500 nM $LaCl_3$. UW154 was then added to 50 µM, and absorbance at $\lambda_{abs}$ = 520 nm was monitored for 15 min.

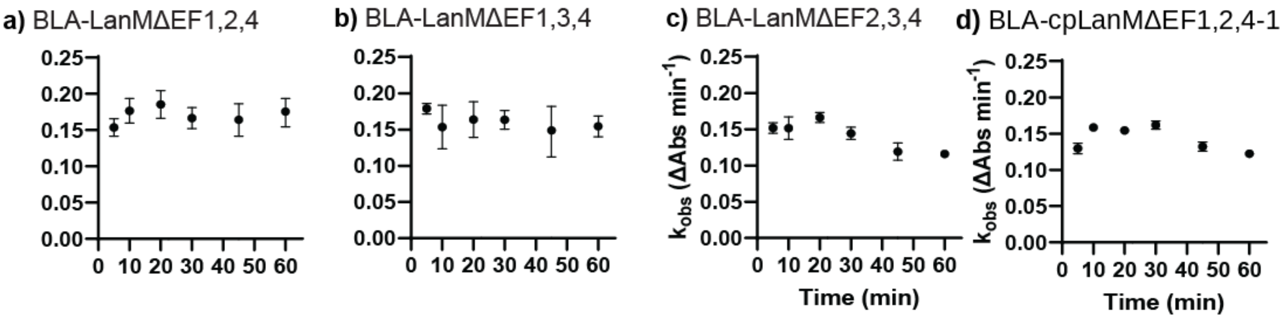


**Supplementary Figure 3.** Response times of the most promising single-binding-site protein switches: a) BLA-LanMΔEF1,2,4, b) BLA-LanMΔEF1,3,4, c) BLA-LanMΔEF2,3,4 and d) BLA-cpLanMΔEF1,2,4-1. Each protein (25 nM) was incubated with 500 nM $LaCl_3$ for 5, 10, 20, 30, 45 or 60 min before UW154 was added. The linear region of each absorbance-time curve was fitted to obtain $k_{obs}$.

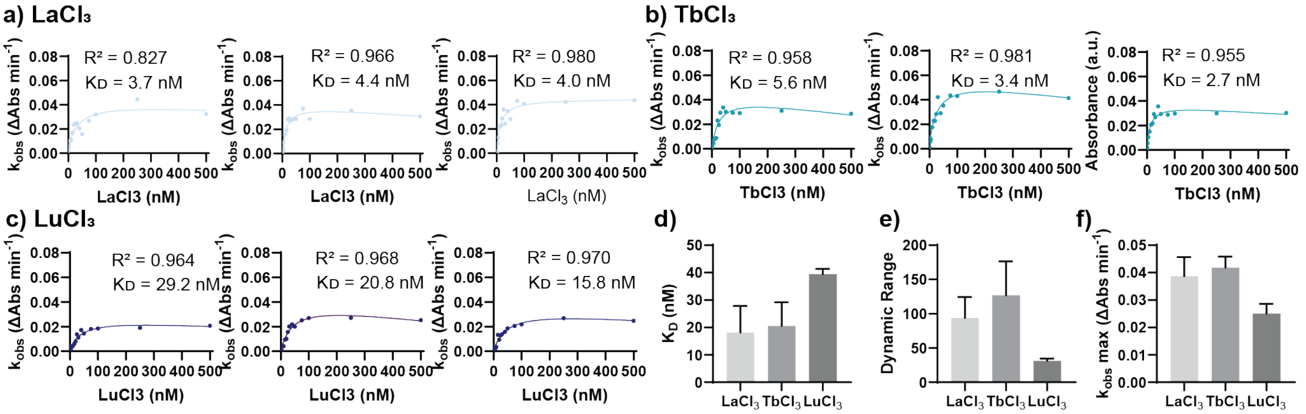


**Supplementary Figure 4.** Apparent affinity of BLA-cpLanMΔEF1,2,4-1 for a) $LaCl_3$, b) $TbCl_3$ and c) $LuCl_3$. Three replicate titrations are shown for each REE. $k_{obs}$ values were obtained from absorbance-time curves recorded after 15 nM BLA-cpLanMΔEF1,2,4-1 was incubated for 10 min with 0–500 nM $LnCl_3$ and then mixed with 50 µM UW154. Curves were fitted to obtain apparent $K_D$ values; the $R^2$ value of each fit is shown. Panels d–f compare, respectively, mean apparent $K_{D,}$ dynamic range and maximum $k_{obs}$ across $LaCl_3$, $TbCl_3$ and $LuCl_3$.

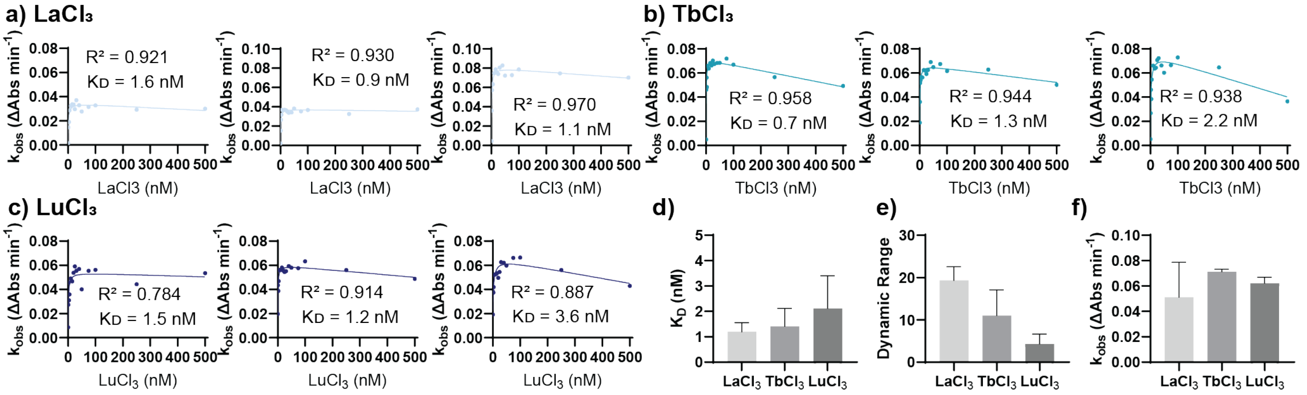


**Supplementary Figure 5.** Apparent affinity of BLA-LanMΔEF2,3,4 for a) $LaCl_3$, b) $TbCl_3$ and c) $LuCl_3$. Three replicate titrations are shown for each REE. $k_{obs}$ values were obtained from absorbance-time curves recorded after 15 nM BLA-LanMΔEF2,3,4 was incubated for 10 min with 0–500 nM $LnCl_3$ and then mixed with 50 µM UW154. Curves were fitted to obtain apparent $K_D$ values; the $R^2$ value of each fit is shown. Panels d–f compare, respectively, mean apparent $K_{D,}$ dynamic range and maximum $k_{obs}$ across $LaCl_3$, $TbCl_3$ and $LuCl_3$.

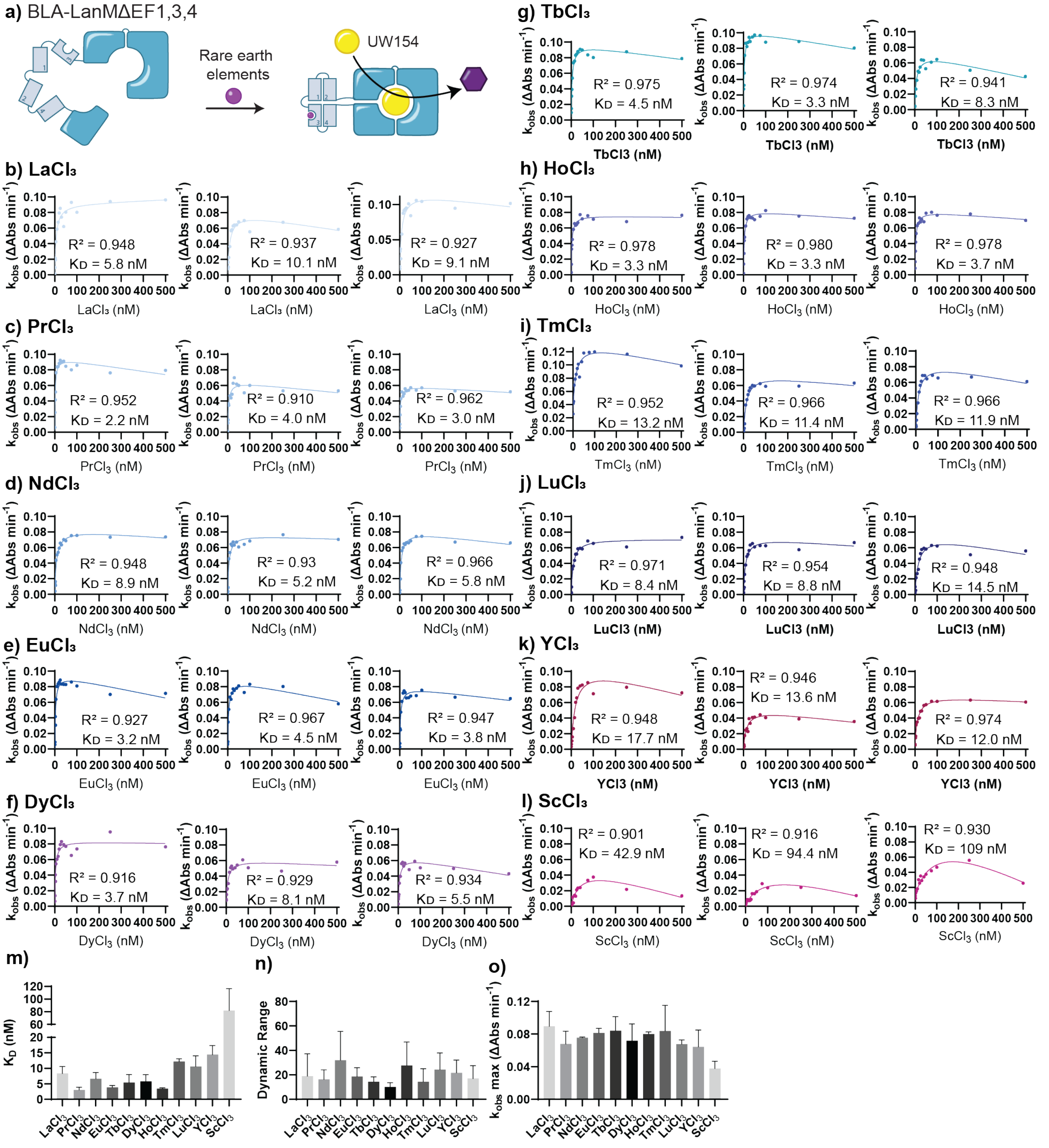


**Supplementary Figure 6.** Apparent affinity of BLA-LanMΔEF1,3,4 for 11 REEs. a) Schematic of REE-dependent activation of the protein switch. b–l) Three replicate titrations for each REE. $k_{obs}$ values were obtained from absorbance-time curves recorded after 15 nM BLA-LanMΔEF1,3,4 was incubated for 10 min with 0–500 nM of the respective REE and then mixed with 50 μM UW154. Curves were fitted to obtain apparent $K_D$ values; the $R^2$ value of each fit is shown. Panels m–o compare, respectively, mean apparent $K_{D,}$ dynamic range and maximum $k_{obs}$ across the REEs.

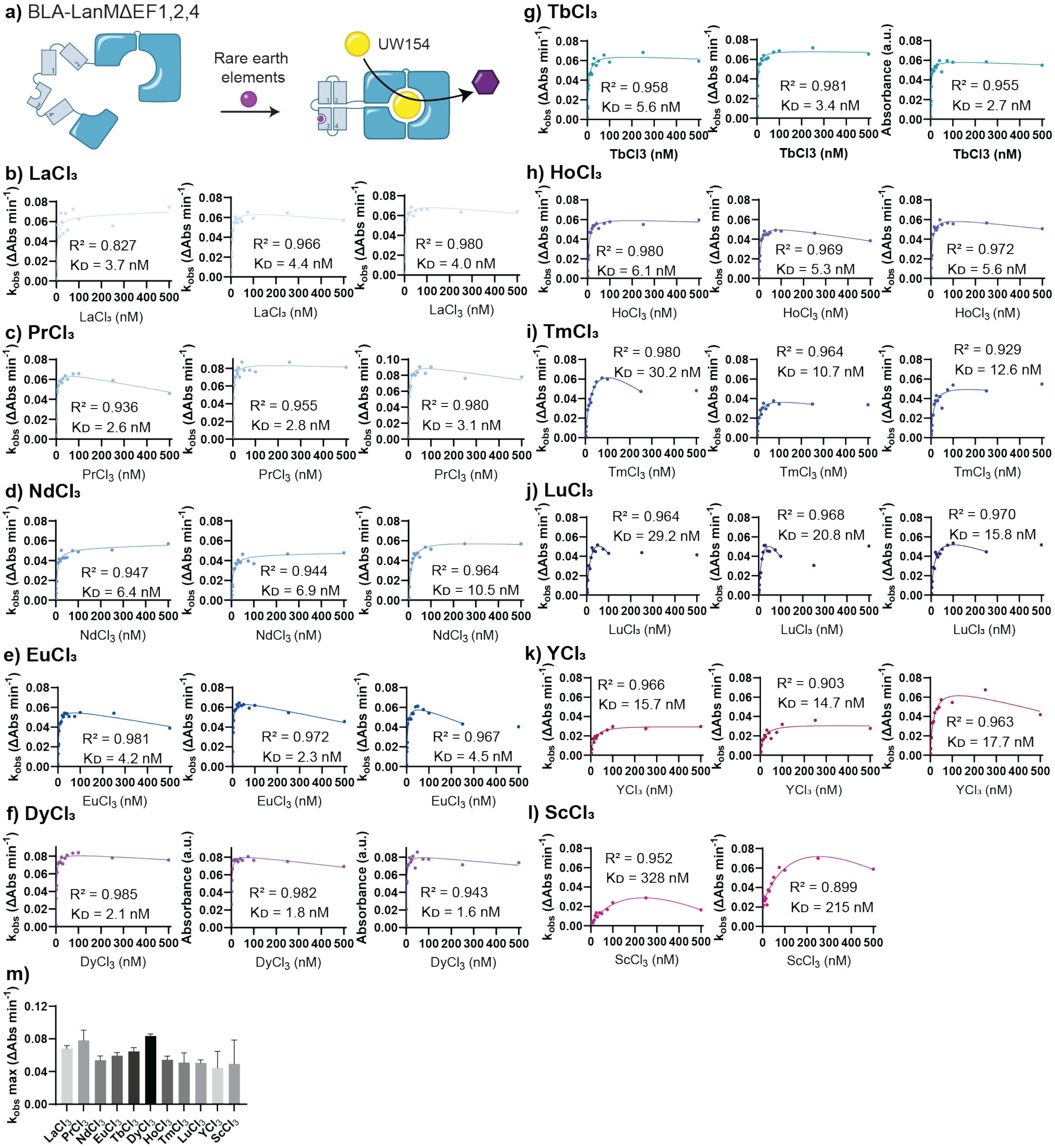

**Supplementary Figure 7.** Apparent affinity of BLA-LanMΔEF1,2,4 for 11 REEs. a) Schematic showing that REE binding activates the protein switch, enabling binding to be monitored from the absorbance of the UW154 reaction product at $\lambda_{abs}$ = 520 nm. b–l) Three replicate titrations for each REE. $k_{obs}$ values were obtained from absorbance-time curves recorded after 15 nM BLA-LanMΔEF1,2,4 was incubated for 10 min with 0–500 nM of the respective REE and then mixed with 50 μM UW154. Curves were fitted to obtain apparent $K_D$ values; the $R^2$ value of each fit is shown. m) Mean maximum catalytic rate ($k_{obs}$,max) for each tested REE.

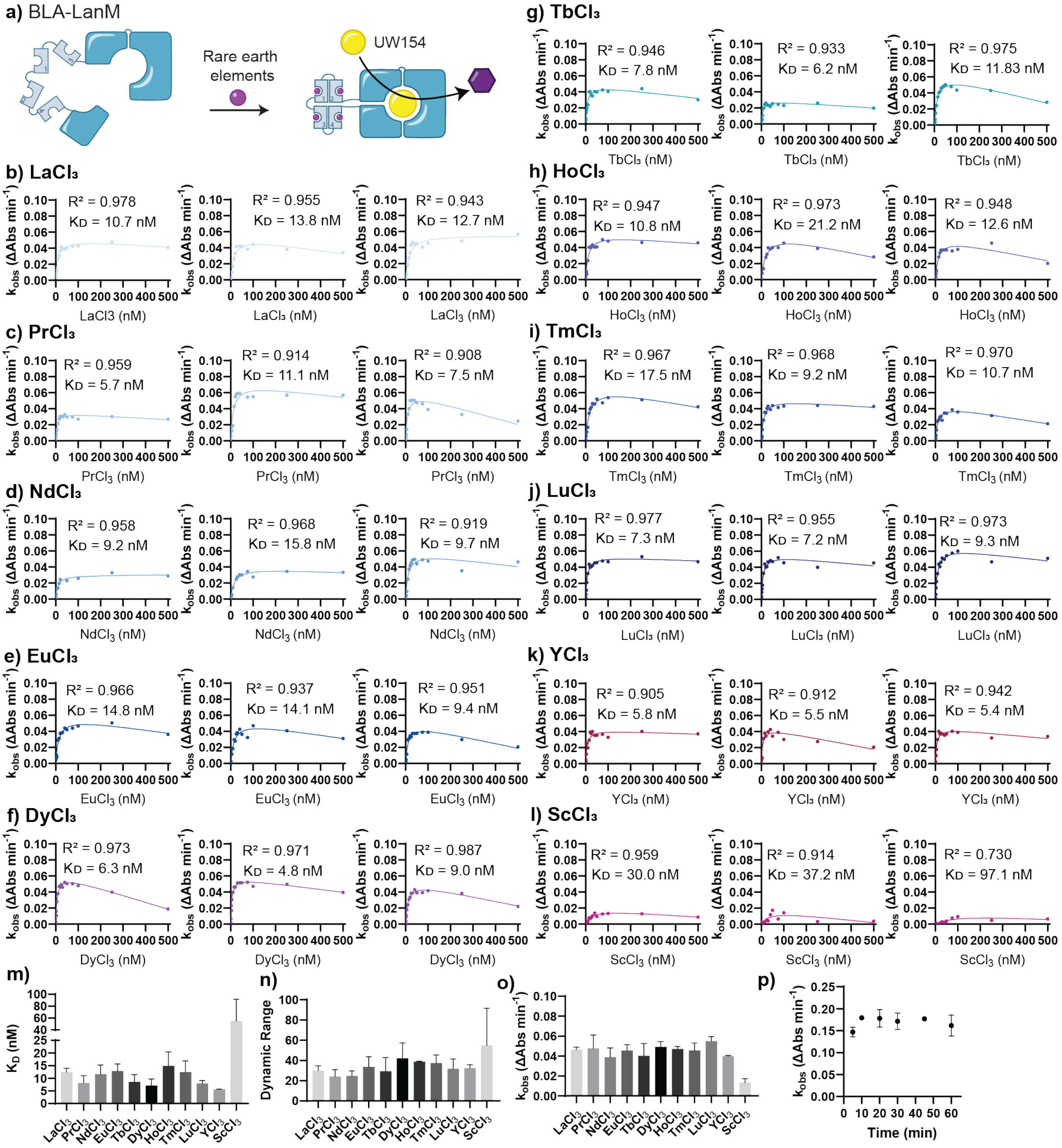


**Supplementary Figure 8.** Apparent affinity of BLA-LanM for 11 REEs. a) Schematic of REE-dependent activation of the protein switch. b–l) Three replicate titrations for each REE. $k_{obs}$ values were obtained from absorbance-time curves recorded after 10 nM BLA-LanM was incubated for 10 min with 0–500 nM of the respective REE and then mixed with 50 µM UW154. Curves were fitted to obtain apparent $K_D$ values; the $R^2$ value of each fit is shown. Panels m–o compare, respectively, mean apparent $K_D$, dynamic range and maximum $k_{obs}$ across the REEs. p) Response time determined by incubating 25 nM BLA-LanM with 500 nM $LaCl_3$ for 5, 10, 20, 30, 45 or 60 min before adding UW154.

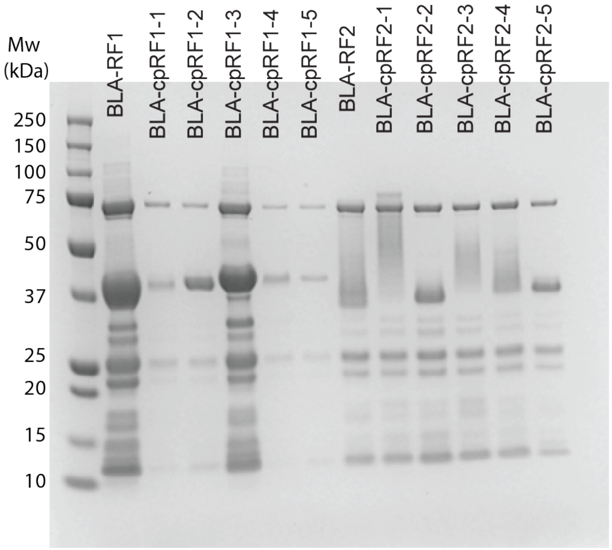


**Supplementary Figure 9.** SDS-PAGE analysis of BLA-RF1 (lane 1), circularly permuted BLA-RF1 variants (lanes 2–6), BLA-RF2 (lane 7) and circularly permuted BLA-RF2 variants (lanes 8–12). Each lane contained 2 μg of protein separated on a 4–12% SDS-PAGE gel and stained with Coomassie Blue.

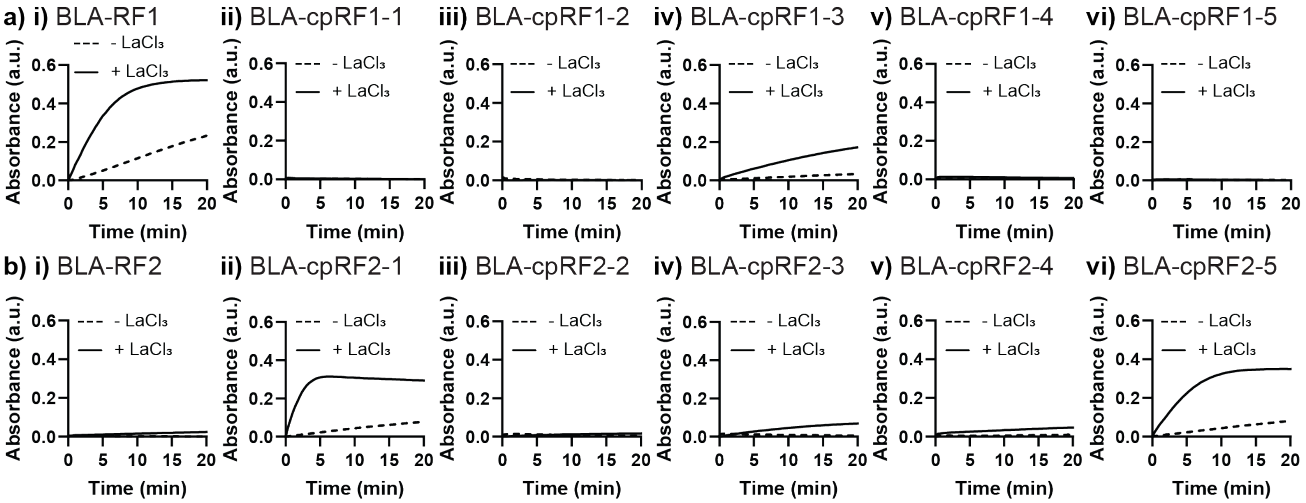


**Supplementary Figure 10.** Activity screening of chimeric BLA switches containing EF-hand-based miniproteins. a) BLA-RF1-derived switches: i) BLA-RF1 and ii–vi) five circularly permuted variants. b) BLA-RF2-derived switches: i) BLA-RF2 and ii–vi) five circularly permuted variants. Purified protein (25 nM) was incubated for 10 min with or without 500 nM $LaCl_3$. UW154 was then added to 50 μM, and absorbance at $\lambda_{abs}$ = 520 nm was monitored for approximately 20 min.

a) BLA-cpRF2-1

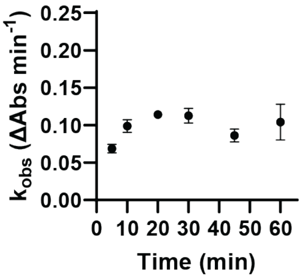


**Supplementary Figure 11.** Response time of BLA-cpRF2-1. Purified BLA-cpRF2-1 (25 nM) was incubated with 500 nM $LaCl_3$ for 5, 10, 20, 30, 45 or 60 min before UW154 was added.

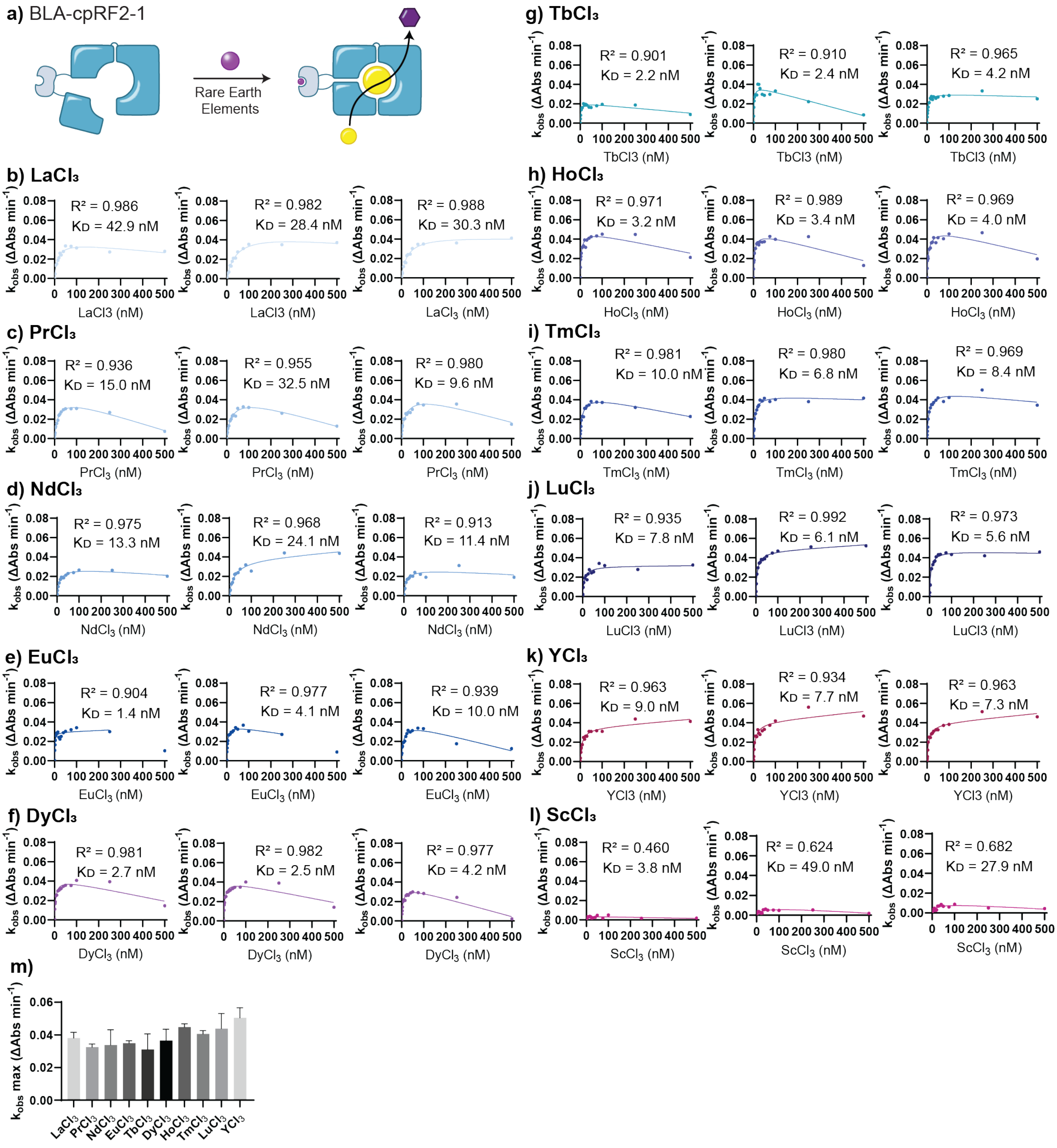


**Supplementary Figure 12.** Apparent affinity of BLA-cpRF2-1 for 11 REEs. a) Schematic of REE-dependent activation of the protein switch, measured from the absorbance of the UW154 reaction product at $\lambda_{abs}$ = 520 nm. b–l) Three replicate titrations for each REE. $k_{obs}$ values were obtained from absorbance-time curves recorded after 10 nM BLA-cpRF2-1 was incubated for 20 min with 0–500 nM of the respective REE and then mixed with 50 μM UW154. Curves were fitted to obtain apparent $K_D$ values; the $R^2$ value of each fit is shown. m) Mean maximum catalytic rate ($k_{obs}$,max) for each tested REE.

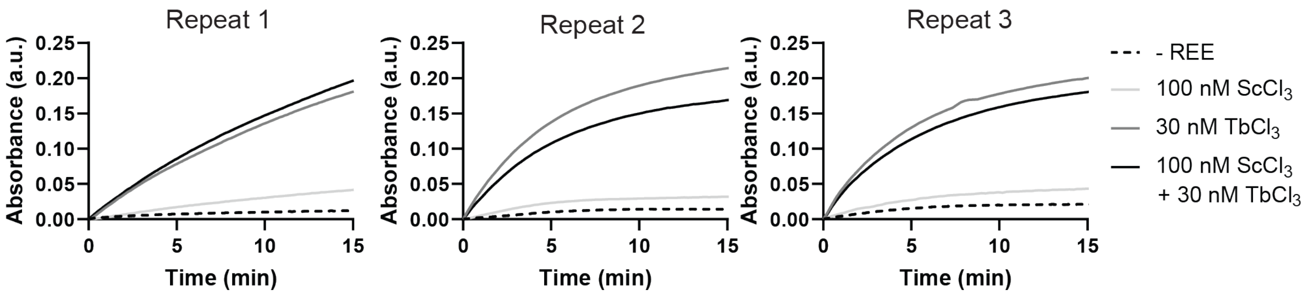


**Supplementary Figure 13.** Three replicate absorbance-time curves for BLA-cpRF2-1 after incubation with no REE, 100 nM $ScCl_3$, 30 nM $TbCl_3$, or 100 nM $ScCl_3$ plus 30 nM $TbCl_3$.

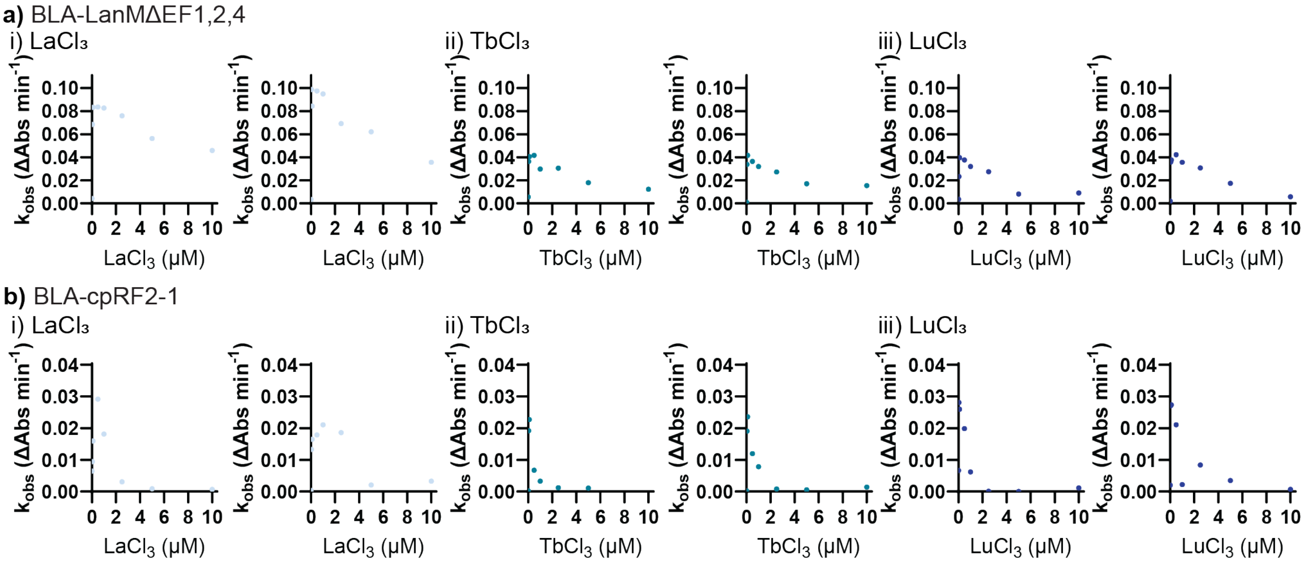


**Supplementary Figure 14.** Concentration-dependent hook effect for a) BLA-LanMΔEF1,2,4 and b) BLA-cpRF2-1. Each switch was incubated with i) $LaCl_3$, ii) $TbCl_3$ or iii) $LuCl_3$ at concentrations up to 10 μM. $k_{obs}$ values were obtained from absorbance-time curves ($\lambda_{abs}$ = 520 nm) after 15 nM BLA-LanMΔEF1,2,4 or 10 nM BLA-cpRF2-1 was incubated with the respective REE for 10 min or 20 min, respectively, and then mixed with 50 μM UW154.

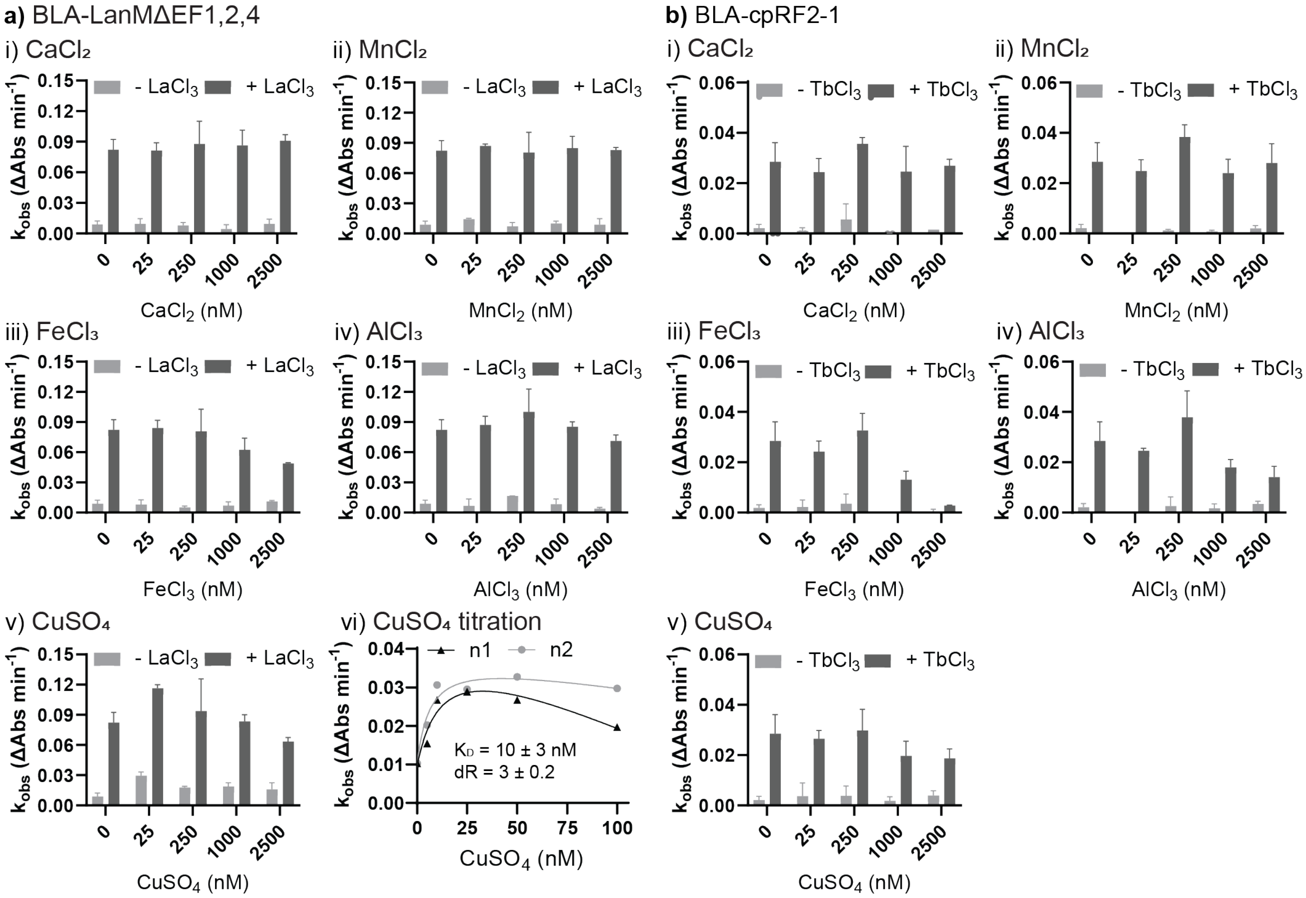


**Supplementary Figure 15.** Selectivity of the purified protein switches for REEs over other metal ions. The switches were incubated with 0, 25, 250, 1,000 or 2,500 nM $CaCl_2$, $MnCl_2$, $FeCl_3$, $AlCl_3$ or $CuSO_4$. a) BLA-LanMΔEF1,2,4 (15 nM) was incubated for 10 min with i) $CaCl_2$, ii) $MnCl_2$, iii) $FeCl_3$, iv) $AlCl_3$ or v) $CuSO_4$, with or without 50 nM $LaCl_3$. $k_{obs}$ values were obtained by fitting the linear region of absorbance-time curves ($\lambda_{abs}$ = 520 nm) after addition of 50 μM UW154. vi) BLA-LanMΔEF1,2,4 (15 nM) was incubated with 0–100 nM $CuSO_4$; $k_{obs}$ values were plotted against $CuSO_4$ concentration, and the curves were fitted to obtain apparent $K_D$ values. The mean dynamic range of two replicates is shown below the curve. b) BLA-cpRF2-1 (10 nM) was incubated for 20 min with i) $CaCl_2$, ii) $MnCl_2$, iii) $FeCl_3$, iv) $AlCl_3$ or v) $CuSO_4$, with or without 30 nM $TbCl_3$. $k_{obs}$ values were obtained as described for a).

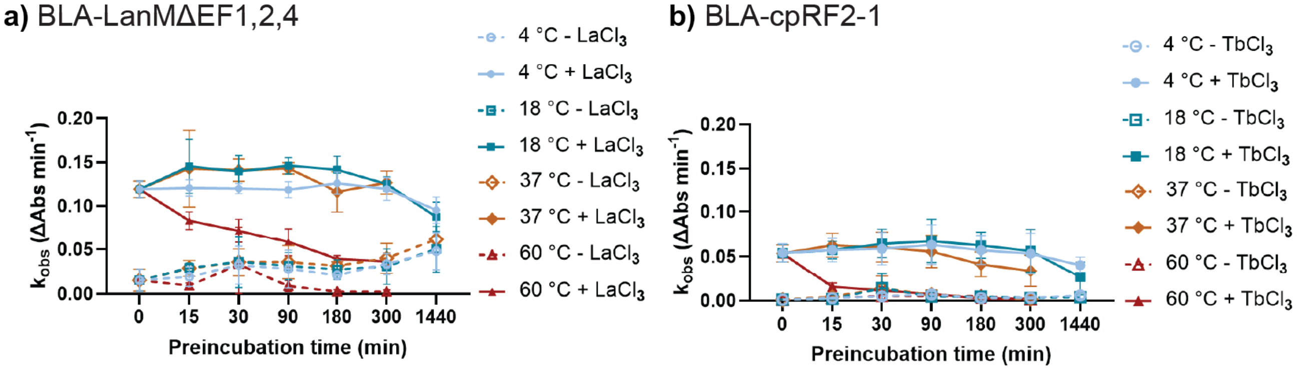


**Supplementary Figure 16.** Thermostability of a) BLA-LanMΔEF1,2,4 and b) BLA-cpRF2-1. Each switch (4 μM) was preincubated at 4, 18, 37 or 60 °C for 0, 15, 30, 90, 180, 300 or 1,440 min. Aliquots were then diluted to 25 nM and incubated for 15 min with a) 50 nM $LaCl_3$ or b) 30 nM $TbCl_3$ before addition of 50 μM

UW154. $k_{obs}$ values were obtained by fitting the linear region of the absorbance-time curves ($\lambda_{abs}$ = 520 nm).

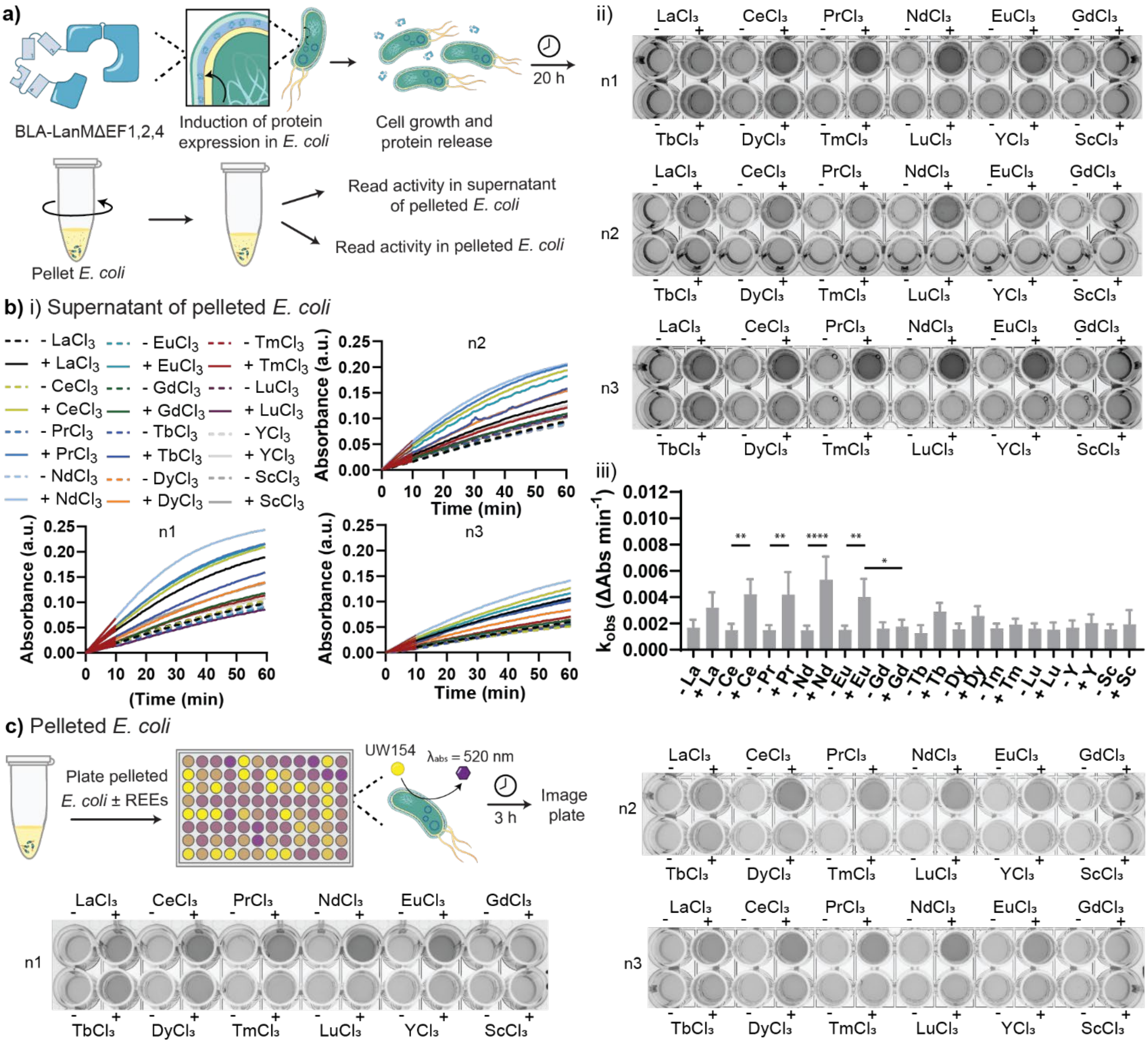


**Supplementary Figure 17.** Activity of *E. coli* expressing BLA-LanMΔEF1,2,4. a) Workflow for periplasmic expression, 20 h cell growth and separation of supernatant and pellet. b) Supernatant assay. Supernatant was incubated for 1 h with 500 nM REE in 20 mM Tris-HCl (pH 7.2), 100 mM NaCl, then mixed with 50 µM UW154. i) Absorbance-time curves recorded for approximately 60 min. ii) End-point ChemiDoc images. iii) $k_{obs}$ values from the linear curve regions compared across 12 REEs. *$p < 0.05$, **$p < 0.005$, ***$p < 0.0005$ and ****$p < 0.00005$; no stars, not significant. c) Pellet assay performed as in b), with imaging 3 h after UW154 addition. Three independent replicates are shown.

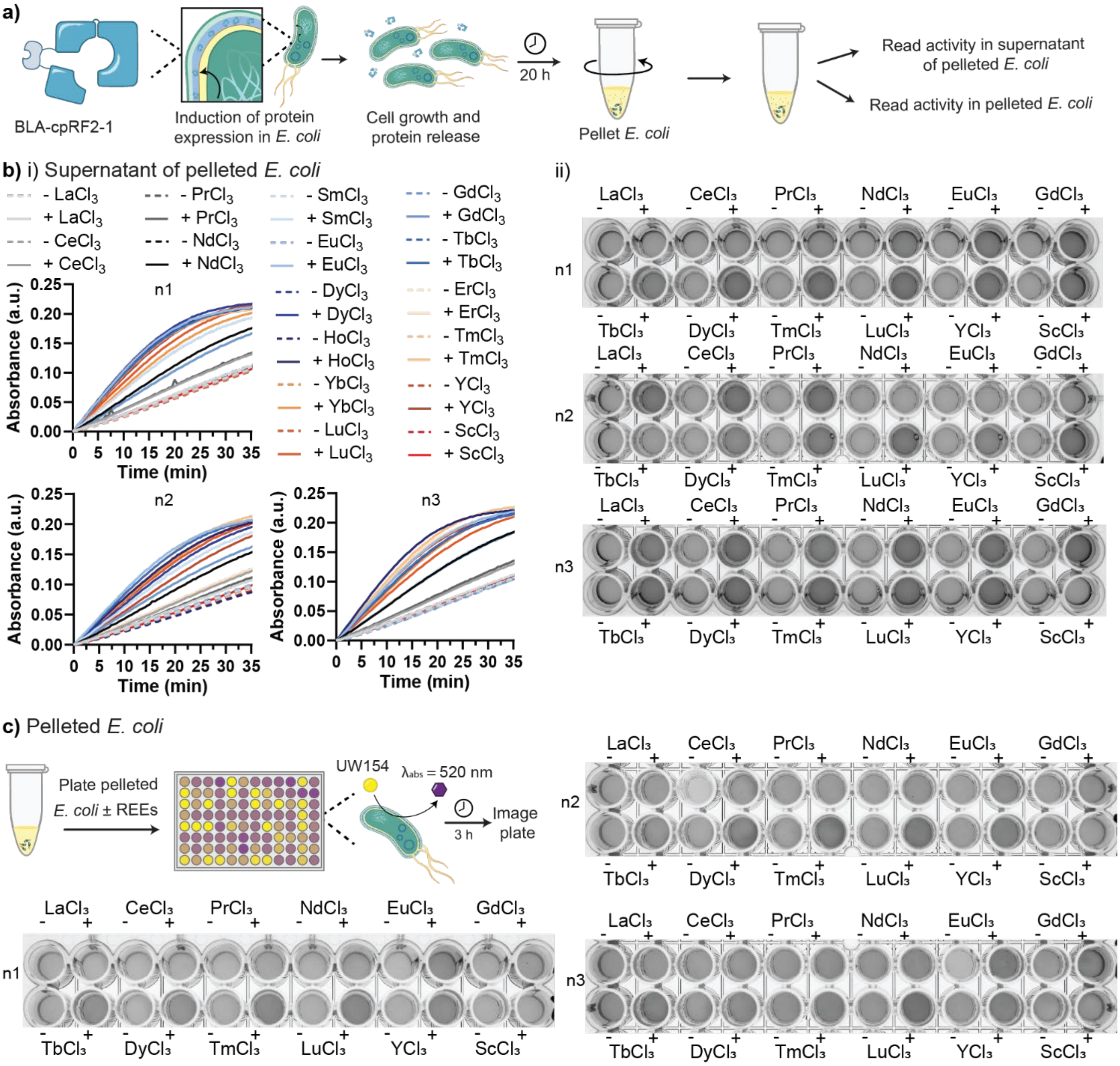


**Supplementary Figure 18.** Activity of *E. coli* expressing BLA-cpRF2-1. a) Workflow for induction of periplasmic expression, cell growth and separation of the supernatant from the cell pellet. b) Supernatant assay. The supernatant was incubated for 1 h with 500 nM REE in 20 mM Tris-HCl (pH 7.2), 100 mM NaCl, and UW154 was then added to 50 μM. i) Absorbance-time curves recorded for approximately 60 min. ii) End-point ChemiDoc images of the 96-well plates. c) Pellet assay. Resuspended cells were incubated with 500 nM REE in the same buffer for 1 h, mixed with 50 μM UW154 and imaged after 3 h. Three independent replicates are shown.

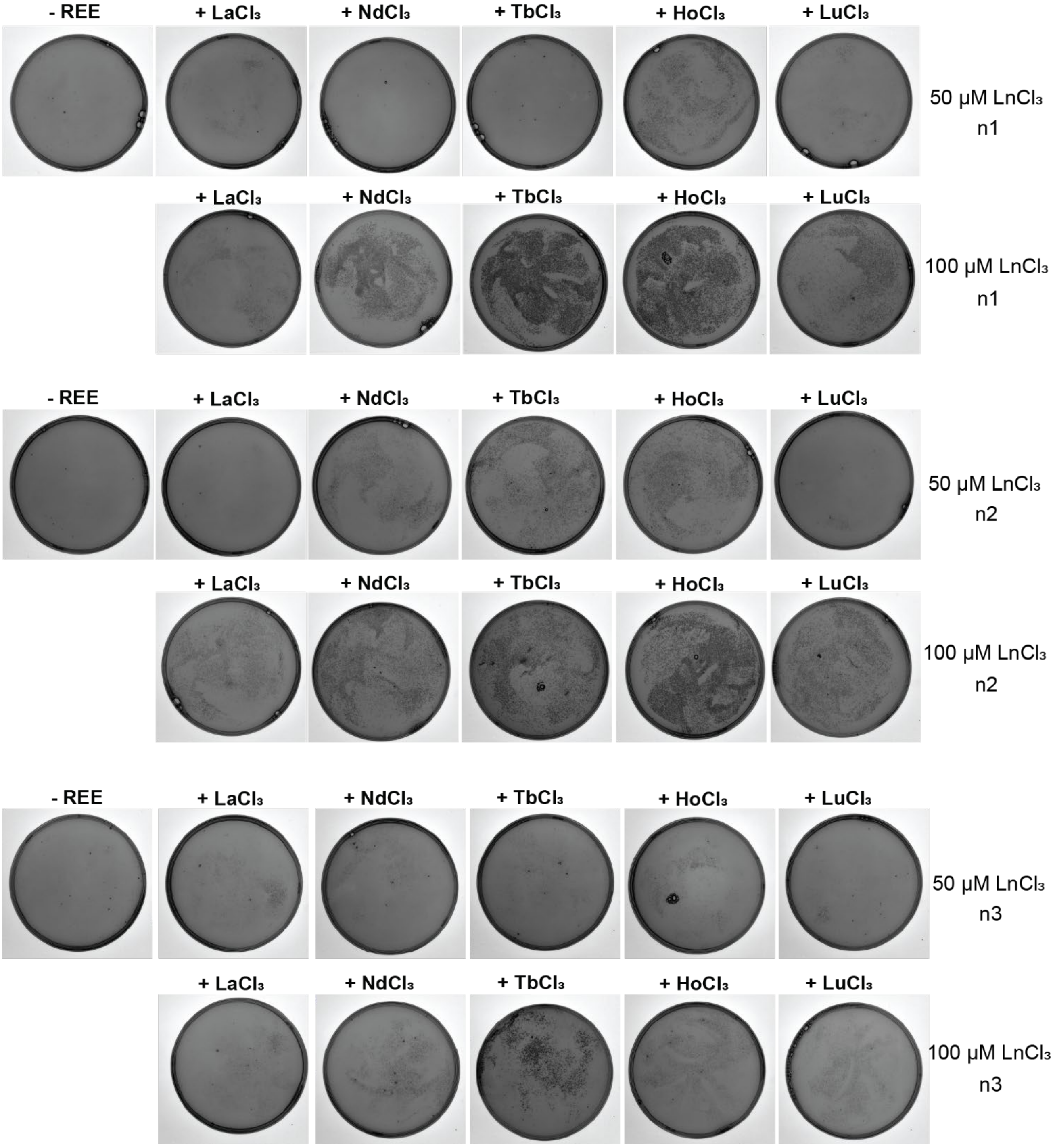


**Supplementary Figure 19.** Autoinduction agar plates containing 50 µg $mL^{-1}$ kanamycin and 50 µg $mL^{-1}$ ampicillin, seeded with *E. coli* expressing BLA-cpRF2-1. Plates were cast without REEs or with 50 or 100 µM of the indicated $LnCl_3$. Three independent replicates (n1–n3) are shown.

**Supplementary Table 1.** Performance parameters of selected constructs. Apparent $K_D$, dynamic range and maximum catalytic rate ($k_{obs,}$max) were determined from 0–500 nM REE titrations using 10 nM BLA-cpRF2-1 or BLA-LanM and 15 nM of the other switches. The $ScCl_3$ $K_D$ for BLA-cpRF2-1 is omitted because $R^2 < 0.7$. Values are from three independent replicates.

| Construct | Parameter | $LaCl_3$ | $PrCl_3$ | $NdCl_3$ | $EuCl_3$ | $TbCl_3$ | $DyCl_3$ | $HoCl_3$ | $TmCl_3$ | $LuCl_3$ | $YCl_3$ | $ScCl_3$ |
|---|---|---|---|---|---|---|---|---|---|---|---|---|
| BLA-LanMΔEF1,2,4 | $K_D$ (nM) | 4.0 ± 0.3 | 2.8 ± 0.2 | 8 ± 2 | 4 ± 1 | 4 ± 1 | 1.9 ± 0.2 | 5.6 ± 0.3 | 18 ± 9 | 22 ± 6 | 16 ± 1 | 272 ± 57 |
| | Dynamic range | 27 ± 2 | 28 ± 7 | 34 ± 3 | 30 ± 5 | 30 ± 2 | 31 ± 8 | 25 ± 2 | 24 ± 1 | 30 ± 8 | 29 ± 4 | 7 ± 3 |
| | $k_{obs,}$max (ΔAbs $min^{-1}$) | 0.068 ± 0.003 | 0.08 ± 0.01 | 0.054 ± 0.004 | 0.059 ± 0.003 | 0.065 ± 0.004 | 0.083 ± 0.002 | 0.054 ± 0.004 | 0.05 ± 0.01 | 0.051 ± 0.003 | 0.04 ± 0.02 | 0.05 ± 0.02 |
| BLA-LanMΔEF1,3,4 | $K_D$ (nM) | 8 ± 2 | 3.1 ± 0.7 | 7 ± 2 | 3.9 ± 0.5 | 5 ± 2 | 6 ± 2 | 3.4 ± 0.2 | 14 ± 3 | 11 ± 3 | 14 ± 2 | 82 ± 28 |
| | Dynamic range | 19 ± 15 | 16 ± 6 | 32 ± 19 | 19 ± 6 | 14 ± 3 | 10 ± 3 | 28 ± 16 | 14 ± 9 | 24 ± 11 | 22 ± 9 | 17 ± 8 |
| | $k_{obs,}$max (ΔAbs $min^{-1}$) | 0.09 ± 0.01 | 0.07 ± 0.01 | 0.075 ± 0.001 | 0.082 ± 0.004 | 0.08 ± 0.01 | 0.07 ± 0.02 | 0.080 ± 0.002 | 0.08 ± 0.03 | 0.068 ± 0.004 | 0.06 ± 0.02 | 0.04 ± 0.01 |
| BLA-LanMΔEF2,3,4 | $K_D$ (nM) | 1.2 ± 0.3 | | | | 1.4 ± 0.6 | | | | 2 ± 1 | | |
| | Dynamic range | 19 ± 2 | | | | 11 ± 5 | | | | 4 ± 2 | | |
| | $k_{obs,}$max (ΔAbs $min^{-1}$) | 0.05 ± 0.02 | | | | 0.071 ± 0.002 | | | | 0.062 ± 0.004 | | |
| BLA-LanM | $K_D$ (nM) | 12 ± 1 | 8 ± 2 | 12 ± 3 | 13 ± 2 | 9 ± 2 | 7 ± 2 | 15 ± 5 | 12 ± 4 | 8 ± 1 | 5.5 ± 0.2 | 55 ± 2 |
| | Dynamic range | 30 ± 4 | 24 ± 6 | 25 ± 4 | 34 ± 8 | 29 ± 11 | 42 ± 12 | 47 ± 12 | 38 ± 7 | 32 ± 8 | 33 ± 3 | 13 ± 7 |
| | $k_{obs,}$max (ΔAbs $min^{-1}$) | 0.047 ± 0.002 | 0.05 ± 0.01 | 0.04 ± 0.01 | 0.045 ± 0.005 | 0.04 ± 0.01 | 0.049 ± 0.004 | 0.047 ± 0.002 | 0.05 ± 0.01 | 0.055 ± 0.004 | 0.040 ± 0.001 | 0.013 ± 0.003 |
| BLA-cpLanMΔEF1,2,4-1 | $K_D$ (nM) | 18 ± 8 | | | | 20 ± 7 | | | | 39 ± 2 | | |
| | Dynamic range | 94 ± 25 | | | | 127 ± 41 | | | | 31 ± 3 | | |
| | $k_{obs,}$max (ΔAbs $min^{-1}$) | 0.042 ± 0.003 | | | | 0.039 ± 0.006 | | | | 0.025 ± 0.003 | | |
| BLA-cpRF2-1 | $K_D$ (nM) | 29 ± 10 | 19 ± 10 | 16 ± 6 | 6 ± 3 | 3 ± 1 | 3.1 ± 0.8 | 3.5 ± 0.3 | 8 ± 1 | 6 ± 1 | 8.0 ± 0.7 | - |
| | Dynamic range | 86 ± 16 | 83 ± 9 | 73 ± 7 | 77 ± 10 | 91 ± 9 | 90 ± 3 | 119 ± 6 | 110 ± 17 | 108 ± 8 | 129 ± 16 | 11 ± 3 |
| | $k_{obs,}$max (ΔAbs $min^{-1}$) | 0.036 ± 0.003 | 0.033 ± 0.002 | 0.034 ± 0.008 | 0.035 ± 0.001 | 0.03 ± 0.01 | 0.036 ± 0.006 | 0.045 ± 0.002 | 0.041 ± 0.002 | 0.044 ± 0.007 | 0.050 ± 0.005 | 0.007 ± 0.001 |

**Supplementary Table 2.** Amino acid sequences of the constructs used in this study.

| Construct | Sequence |
|---|---|
| BLA-LanMΔEF1,2,4 | DHPETLVKVKDAEDQLGSGVDIAAFDPSKSGTIDLKEALAAGSAAFDKLDP SKSGTLDAKELKGRVSEADLKKLDPDNDGTLDKKEYLAAVEAQFKAANPS NSGTIDARELASPAGSALVNLIRGSGARVGYIELDLNSGKILESFRPEERFP |

| | |
|---|---|
| | MMSTFKVLLCGAVLSRVDAGQEQLGRRIHYSQNDLVEYSPVTEKHLTDGM TVRELCSAAITMSDNTAANLLLTTIGGPKELTAFLHNMGDHVTRLDRWEPE LNEAIPNDERDTTMPVAMATTLRKLLTGELLTLASRQQLIDWMEADKVAGP LLRSALPAGWFIADKSGAGERGSRGIIAALGPDGKPSRIVVIYTTGSQATMD ERNRQIAEIGASLIKHWGKLAAALEHHHHHH |
| BLA-LanMΔEF1,3,4 | DHPETLVKVKDAEDQLGSGVDIAAFDPSKSGTIDLKEALAAGSAAFDKLDP DKDGTLDAKELKGRVSEADLKKLDPSNSGTLDKKEYLAAVEAQFKAANPS NSGTIDARELASPAGSALVNLIRGSGARVGYIELDLNSGKILESFRPEERFP MMSTFKVLLCGAVLSRVDAGQEQLGRRIHYSQNDLVEYSPVTEKHLTDGM TVRELCSAAITMSDNTAANLLLTTIGGPKELTAFLHNMGDHVTRLDRWEPE LNEAIPNDERDTTMPVAMATTLRKLLTGELLTLASRQQLIDWMEADKVAGP LLRSALPAGWFIADKSGAGERGSRGIIAALGPDGKPSRIVVIYTTGSQATMD ERNRQIAEIGASLIKHWGKLAAALEHHHHHH |
| BLA-LanMΔEF2,3,4 | DHPETLVKVKDAEDQLGARVGYIELDLNSGKILESFRPEERFPMMSTFKVLL CGAVLSRVDAGQEQLGRRIHYSQNDLVEYSPVTEKHLTDGMTVRELCSAAI TMSDNTAANLLLTTIGGPKELTAFLHNMGDHVTRLDRWEPELNEAIPNDER DTTMPVAMATTLRKLLTGELLTLASRQQLIDWMEADKVAGPLLRSALPAGW FIADKSGAGERGSRGIIAALGPDGSGVDIAAFDPDKDGTIDLKEALAAGSAA FDKLDPSKSGTLDAKELKGRVSEADLKKLDPSNSGTLDKKEYLAAVEAQFK AANPSNSGTIDARELASPAGSALVNLIRGSGKPSRIVVIYTTGSQATMDERN RQIAEIGASLIKHWGKLAAALEHHHHHH |
| BLA-LanM | DHPETLVKVKDAEDQLGSGVDIAAFDPDKDGTIDLKEALAAGSAAFDKLDP DKDGTLDAKELKGRVSEADLKKLDPDNDGTLDKKEYLAAVEAQFKAANPD NDGTIDARELASPAGSALVNLIRGSGARVGYIELDLNSGKILESFRPEERFP MMSTFKVLLCGAVLSRVDAGQEQLGRRIHYSQNDLVEYSPVTEKHLTDGM TVRELCSAAITMSDNTAANLLLTTIGGPKELTAFLHNMGDHVTRLDRWEPE LNEAIPNDERDTTMPVAMATTLRKLLTGELLTLASRQQLIDWMEADKVAGP LLRSALPAGWFIADKSGAGERGSRGIIAALGPDGKPSRIVVIYTTGSQATMD ERNRQIAEIGASLIKHWGKLAAALEHHHHHH |
| BLA-cpLanMΔEF1,2,4-1 | DHPETLVKVKDAEDQLGKSGTLDAKELKGRVSEADLKKLDPDNDGTLDKK EYLAAVEAQFKAANPSNSGTIDARELASPAGSALVNLIRGGGSGGSGGGP TTTTKVDIAAFDPSKSGTIDLKEALAAGSAAFDKLDPGARVGYIELDLNSGKI LESFRPEERFPMMSTFKVLLCGAVLSRVDAGQEQLGRRIHYSQNDLVEYSP VTEKHLTDGMTVRELCSAAITMSDNTAANLLLTTIGGPKELTAFLHNMGDH VTRLDRWEPELNEAIPNDERDTTMPVAMATTLRKLLTGELLTLASRQQLID WMEADKVAGPLLRSALPAGWFIADKSGAGERGSRGIIAALGPDGKPSRIVV IYTTGSQATMDERNRQIAEIGASLIKHWGKLAAALEHHHHHH |
| BLA-cpLanMΔEF1,2,4-2 | DHPETLVKVKDAEDQLGADLKKLDPDNDGTLDKKEYLAAVEAQFKAANPS NSGTIDARELASPAGSALVNLIRGGGSGGSGGGPTTTTKVDIAAFDPSKSG TIDLKEALAAGSAAFDKLDPSKSGTLDAKELKGRVSGARVGYIELDLNSGKI LESFRPEERFPMMSTFKVLLCGAVLSRVDAGQEQLGRRIHYSQNDLVEYSP VTEKHLTDGMTVRELCSAAITMSDNTAANLLLTTIGGPKELTAFLHNMGDH VTRLDRWEPELNEAIPNDERDTTMPVAMATTLRKLLTGELLTLASRQQLID WMEADKVAGPLLRSALPAGWFIADKSGAGERGSRGIIAALGPDGKPSRIVV IYTTGSQATMDERNRQIAEIGASLIKHWGKLAAALEHHHHHH |
| BLA-cpLanMΔEF1,2,4-3 | DHPETLVKVKDAEDQLGGTLDKKEYLAAVEAQFKAANPSNSGTIDARELAS PAGSALVNLIRGGGSGGSGGGPTTTTKVDIAAFDPSKSGTIDLKEALAAGSA AFDKLDPSKSGTLDAKELKGRVSEADLKKLDPDNGARVGYIELDLNSGKIL ESFRPEERFPMMSTFKVLLCGAVLSRVDAGQEQLGRRIHYSQNDLVEYSP VTEKHLTDGMTVRELCSAAITMSDNTAANLLLTTIGGPKELTAFLHNMGDH VTRLDRWEPELNEAIPNDERDTTMPVAMATTLRKLLTGELLTLASRQQLID |

| | |
|---|---|
| | WMEADKVAGPLLRSALPAGWFIADKSGAGERGSRGIIAALGPDGKPSRIVV<br>IYTTGSQATMDERNRQIAEIGASLIKHWGKLAAALEHHHHHH |
| BLA-cpLanMΔEF1,2,4-4 | DHPETLVKVKDAEDQLGGTIDARELASPAGSALVNLIRGGGSGGSGGGPT<br>TTTKVDIAAFDPSKSGTIDLKEALAAGSAAFDKLDPSKSGTLDAKELKGRVSE<br>ADLKKLDPDNDGTLDKKEYLAAVEAQFKAANPSNGARVGYIELDLNSGKIL<br>ESFRPEERFPMMSTFKVLLCGAVLSRVDAGQEQLGRRIHYSQNDLVEYSP<br>VTEKHLTDGMTVRELCSAAITMSDNTAANLLLTTIGGPKELTAFLHNMGDH<br>VTRLDRWEPELNEAIPNDERDTTMPVAMATTLRKLLTGELLTLASRQQLID<br>WMEADKVAGPLLRSALPAGWFIADKSGAGERGSRGIIAALGPDGKPSRIVV<br>IYTTGSQATMDERNRQIAEIGASLIKHWGKLAAALEHHHHHH |
| BLA-cpLanMΔEF1,2,4-5 | DHPETLVKVKDAEDQLGAGSALVNLIRGGGSGGSGGGPTTTTKVDIAAFD<br>PSKSGTIDLKEALAAGSAAFDKLDPSKSGTLDAKELKGRVSEADLKKLDPD<br>NDGTLDKKEYLAAVEAQFKAANPSNSGTIDARELASGARVGYIELDLNSGK<br>ILESFRPEERFPMMSTFKVLLCGAVLSRVDAGQEQLGRRIHYSQNDLVEYS<br>PVTEKHLTDGMTVRELCSAAITMSDNTAANLLLTTIGGPKELTAFLHNMGD<br>HVTRLDRWEPELNEAIPNDERDTTMPVAMATTLRKLLTGELLTLASRQQLID<br>WMEADKVAGPLLRSALPAGWFIADKSGAGERGSRGIIAALGPDGKPSRIVV<br>IYTTGSQATMDERNRQIAEIGASLIKHWGKLAAALEHHHHHH |
| BLA-cpLanMΔEF1,3,4-1 | DHPETLVKVKDAEDQLGKDGTLDAKELKGRVSEADLKKLDPSNSGTLDKK<br>EYLAAVEAQFKAANPSNSGTIDARELASPAGSALVNLIRGGGSGGSGGGP<br>TTTTKVDIAAFDPSKSGTIDLKEALAAGSAAFDKLDPGARVGYIELDLNSGKI<br>LESFRPEERFPMMSTFKVLLCGAVLSRVDAGQEQLGRRIHYSQNDLVEYSP<br>VTEKHLTDGMTVRELCSAAITMSDNTAANLLLTTIGGPKELTAFLHNMGDH<br>VTRLDRWEPELNEAIPNDERDTTMPVAMATTLRKLLTGELLTLASRQQLID<br>WMEADKVAGPLLRSALPAGWFIADKSGAGERGSRGIIAALGPDGKPSRIVV<br>IYTTGSQATMDERNRQIAEIGASLIKHWGKLAAALEHHHHHH |
| BLA-cpLanMΔEF1,3,4-2 | DHPETLVKVKDAEDQLGADLKKLDPSNSGTLDKKEYLAAVEAQFKAANPS<br>NSGTIDARELASPAGSALVNLIRGGGSGGSGGGPTTTTKVDIAAFDPSKSG<br>TIDLKEALAAGSAAFDKLDPDKDGTLDAKELKGRVSGARVGYIELDLNSGKI<br>LESFRPEERFPMMSTFKVLLCGAVLSRVDAGQEQLGRRIHYSQNDLVEYSP<br>VTEKHLTDGMTVRELCSAAITMSDNTAANLLLTTIGGPKELTAFLHNMGDH<br>VTRLDRWEPELNEAIPNDERDTTMPVAMATTLRKLLTGELLTLASRQQLID<br>WMEADKVAGPLLRSALPAGWFIADKSGAGERGSRGIIAALGPDGKPSRIVV<br>IYTTGSQATMDERNRQIAEIGASLIKHWGKLAAALEHHHHHH |
| BLA-cpLanMΔEF1,3,4-3 | DHPETLVKVKDAEDQLGGTLDKKEYLAAVEAQFKAANPSNSGTIDARELAS<br>PAGSALVNLIRGGGSGGSGGGPTTTTKVDIAAFDPSKSGTIDLKEALAAGSA<br>AFDKLDPDKDGTLDAKELKGRVSEADLKKLDPSNGARVGYIELDLNSGKIL<br>ESFRPEERFPMMSTFKVLLCGAVLSRVDAGQEQLGRRIHYSQNDLVEYSP<br>VTEKHLTDGMTVRELCSAAITMSDNTAANLLLTTIGGPKELTAFLHNMGDH<br>VTRLDRWEPELNEAIPNDERDTTMPVAMATTLRKLLTGELLTLASRQQLID<br>WMEADKVAGPLLRSALPAGWFIADKSGAGERGSRGIIAALGPDGKPSRIVV<br>IYTTGSQATMDERNRQIAEIGASLIKHWGKLAAALEHHHHHH |
| BLA-cpLanMΔEF1,3,4-4 | DHPETLVKVKDAEDQLGGTIDARELASPAGSALVNLIRGGGSGGSGGGPT<br>TTTKVDIAAFDPSKSGTIDLKEALAAGSAAFDKLDPDKDGTLDAKELKGRVS<br>EADLKKLDPSNSGTLDKKEYLAAVEAQFKAANPSNGARVGYIELDLNSGKI<br>LESFRPEERFPMMSTFKVLLCGAVLSRVDAGQEQLGRRIHYSQNDLVEYSP<br>VTEKHLTDGMTVRELCSAAITMSDNTAANLLLTTIGGPKELTAFLHNMGDH<br>VTRLDRWEPELNEAIPNDERDTTMPVAMATTLRKLLTGELLTLASRQQLID<br>WMEADKVAGPLLRSALPAGWFIADKSGAGERGSRGIIAALGPDGKPSRIVV<br>IYTTGSQATMDERNRQIAEIGASLIKHWGKLAAALEHHHHHH |

| | |
|---|---|
| BLA-cpLanMΔEF1,3,4-5 | DHPETLVKVKDAEDQLGAGSALVNLIRGGGSGGSGGGPTTTTKVDIAAFD PSKSGTIDLKEALAAGSAAFDKLDPDKDGTLDAKELKGRVSEADLKKLDPS NSGTLDKKEYLAAVEAQFKAANPSNSGTIDARELASGARVGYIELDLNSGKI LESFRPEERFPMMSTFKVLLCGAVLSRVDAGQEQLGRRIHYSQNDLVEYSP VTEKHLTDGMTVRELCSAAITMSDNTAANLLLTTIGGPKELTAFLHNMGDH VTRLDRWEPELNEAIPNDERDTTMPVAMATTLRKLLTGELLTLASRQQLID WMEADKVAGPLLRSALPAGWFIADKSGAGERGSRGIIAALGPDGKPSRIVV IYTTGSQATMDERNRQIAEIGASLIKHWGKLAAALEHHHHHH |
| BLA-cpLanMΔEF2,3,4-1 | DHPETLVKVKDAEDQLGKSGTLDAKELKGRVSEADLKKLDPSNSGTLDKKE YLAAVEAQFKAANPSNSGTIDARELASPAGSALVNLIRGGGSGGSGGGPTT TTKVDIAAFDPDKDGTIDLKEALAAGSAAFDKLDPGARVGYIELDLNSGKIL ESFRPEERFPMMSTFKVLLCGAVLSRVDAGQEQLGRRIHYSQNDLVEYSP VTEKHLTDGMTVRELCSAAITMSDNTAANLLLTTIGGPKELTAFLHNMGDH VTRLDRWEPELNEAIPNDERDTTMPVAMATTLRKLLTGELLTLASRQQLID WMEADKVAGPLLRSALPAGWFIADKSGAGERGSRGIIAALGPDGKPSRIVV IYTTGSQATMDERNRQIAEIGASLIKHWGKLAAALEHHHHHH |
| BLA-cpLanMΔEF2,3,4-2 | DHPETLVKVKDAEDQLGADLKKLDPSNSGTLDKKEYLAAVEAQFKAANPS NSGTIDARELASPAGSALVNLIRGGGSGGSGGGPTTTTKVDIAAFDPDKDG TIDLKEALAAGSAAFDKLDPSKSGTLDAKELKGRVSGARVGYIELDLNSGKI LESFRPEERFPMMSTFKVLLCGAVLSRVDAGQEQLGRRIHYSQNDLVEYSP VTEKHLTDGMTVRELCSAAITMSDNTAANLLLTTIGGPKELTAFLHNMGDH VTRLDRWEPELNEAIPNDERDTTMPVAMATTLRKLLTGELLTLASRQQLID WMEADKVAGPLLRSALPAGWFIADKSGAGERGSRGIIAALGPDGKPSRIVV IYTTGSQATMDERNRQIAEIGASLIKHWGKLAAALEHHHHHH |
| BLA-cpLanMΔEF2,3,4-3 | DHPETLVKVKDAEDQLGGTLDKKEYLAAVEAQFKAANPSNSGTIDARELAS PAGSALVNLIRGGGSGGSGGGPTTTTKVDIAAFDPDKDGTIDLKEALAAGS AAFDKLDPSKSGTLDAKELKGRVSEADLKKLDPSNGARVGYIELDLNSGKIL ESFRPEERFPMMSTFKVLLCGAVLSRVDAGQEQLGRRIHYSQNDLVEYSP VTEKHLTDGMTVRELCSAAITMSDNTAANLLLTTIGGPKELTAFLHNMGDH VTRLDRWEPELNEAIPNDERDTTMPVAMATTLRKLLTGELLTLASRQQLID WMEADKVAGPLLRSALPAGWFIADKSGAGERGSRGIIAALGPDGKPSRIVV IYTTGSQATMDERNRQIAEIGASLIKHWGKLAAALEHHHHHH |
| BLA-cpLanMΔEF2,3,4-4 | DHPETLVKVKDAEDQLGGTIDARELASPAGSALVNLIRGGGSGGSGGGPT TTTKVDIAAFDPDKDGTIDLKEALAAGSAAFDKLDPSKSGTLDAKELKGRVS EADLKKLDPSNSGTLDKKEYLAAVEAQFKAANPSNGARVGYIELDLNSGKI LESFRPEERFPMMSTFKVLLCGAVLSRVDAGQEQLGRRIHYSQNDLVEYSP VTEKHLTDGMTVRELCSAAITMSDNTAANLLLTTIGGPKELTAFLHNMGDH VTRLDRWEPELNEAIPNDERDTTMPVAMATTLRKLLTGELLTLASRQQLID WMEADKVAGPLLRSALPAGWFIADKSGAGERGSRGIIAALGPDGKPSRIVV IYTTGSQATMDERNRQIAEIGASLIKHWGKLAAALEHHHHHH |
| BLA-cpLanMΔEF2,3,4-5 | DHPETLVKVKDAEDQLGAGSALVNLIRGGGSGGSGGGPTTTTKVDIAAFD PDKDGTIDLKEALAAGSAAFDKLDPSKSGTLDAKELKGRVSEADLKKLDPS NSGTLDKKEYLAAVEAQFKAANPSNSGTIDARELASGARVGYIELDLNSGKI LESFRPEERFPMMSTFKVLLCGAVLSRVDAGQEQLGRRIHYSQNDLVEYSP VTEKHLTDGMTVRELCSAAITMSDNTAANLLLTTIGGPKELTAFLHNMGDH VTRLDRWEPELNEAIPNDERDTTMPVAMATTLRKLLTGELLTLASRQQLID WMEADKVAGPLLRSALPAGWFIADKSGAGERGSRGIIAALGPDGKPSRIVV IYTTGSQATMDERNRQIAEIGASLIKHWGKLAAALEHHHHHH |
| BLA-RF1 | DHPETLVKVKDAEDQLGARVGYIELDLNSGKILESFRPEERFPMMSTFKVLL CGAVLSRVDAGQEQLGRRIHYSQNDLVEYSPVTEKHLTDGMTVRELCSAAI TMSDNTAANLLLTTIGGPKELTAFLHNMGDHVTRLDRWEPELNEAIPNDER |

| | |
|---|---|
| | DTTMPVAMATTLRKLLTGELLTLASRQQLIDWMEADKVAGPLLRSALPAGW<br>FIADKSGAGERGSRGIIAALGPDGGSGGKKVITISEDTPESFEVKMGKGLLK<br>VTVPGKTFIVEDPDKDGWLDAKEIKALLAAAKAATGAKTIEIEDIPKGGSGGK<br>PSRIVVIYTTGSQATMDERNRQIAEIGASLIKHWGKLAAALEHHHHHH |
| BLA-RF2 | DHPETLVKVKDAEDQLGARVGYIELDLNSGKILESFRPEERFPMMSTFKVLL<br>CGAVLSRVDAGQEQLGRRIHYSQNDLVEYSPVTEKHLTDGMTVRELCSAAI<br>TMSDNTAANLLLTTIGGPKELTAFLHNMGDHVTRLDRWEPELNEAIPNDER<br>DTTMPVAMATTLRKLLTGELLTLASRQQLIDWMEADKVAGPLLRSALPAGW<br>FIADKSGAGERGSRGIIAALGPDGSGRRAYLLRVDPDTLVEISPEEAERLAKT<br>RPVLEVEDPDKDGWLDAKERAWILEHLRETHPDASGAVIVVVDGSGKPSR<br>IVVIYTTGSQATMDERNRQIAEIGASLIKHWGKLAAALEHHHHHH |
| BLA-cpRF1-1 | DHPETLVKVKDAEDQLGARVGYIELDLNSGKILESFRPEERFPMMSTFKVLL<br>CGAVLSRVDAGQEQLGRRIHYSQNDLVEYSPVTEKHLTDGMTVRELCSAAI<br>TMSDNTAANLLLTTIGGPKELTAFLHNMGDHVTRLDRWEPELNEAIPNDER<br>DTTMPVAMATTLRKLLTGELLTLASRQQLIDWMEADKVAGPLLRSALPAGW<br>FIADKSGAGERGSRGIIAALGPDGESFEVKMGKGLLKVTVPGKTFIVEDPDK<br>DGWLDAKEIKALLAAAKAATGAKTIEIEDIPKGGSGGSGGKKVITISEDTGKP<br>SRIVVIYTTGSQATMDERNRQIAEIGASLIKHWGKLAAALEHHHHHH |
| BLA-cpRF1-2 | DHPETLVKVKDAEDQLGARVGYIELDLNSGKILESFRPEERFPMMSTFKVLL<br>CGAVLSRVDAGQEQLGRRIHYSQNDLVEYSPVTEKHLTDGMTVRELCSAAI<br>TMSDNTAANLLLTTIGGPKELTAFLHNMGDHVTRLDRWEPELNEAIPNDER<br>DTTMPVAMATTLRKLLTGELLTLASRQQLIDWMEADKVAGPLLRSALPAGW<br>FIADKSGAGERGSRGIIAALGPDGLLKVTVPGKTFIVEDPDKDGWLDAKEIK<br>ALLAAAKAATGAKTIEIEDIPKGGSGGSGGKKVITISEDTPESFEVKMGKGKP<br>SRIVVIYTTGSQATMDERNRQIAEIGASLIKHWGKLAAALEHHHHHH |
| BLA-cpRF1-3 | DHPETLVKVKDAEDQLGARVGYIELDLNSGKILESFRPEERFPMMSTFKVLL<br>CGAVLSRVDAGQEQLGRRIHYSQNDLVEYSPVTEKHLTDGMTVRELCSAAI<br>TMSDNTAANLLLTTIGGPKELTAFLHNMGDHVTRLDRWEPELNEAIPNDER<br>DTTMPVAMATTLRKLLTGELLTLASRQQLIDWMEADKVAGPLLRSALPAGW<br>FIADKSGAGERGSRGIIAALGPDGKTFIVEDPDKDGWLDAKEIKALLAAAKA<br>ATGAKTIEIEDIPKGGSGGSGGKKVITISEDTPESFEVKMGKGLLKVTVPGKP<br>SRIVVIYTTGSQATMDERNRQIAEIGASLIKHWGKLAAALEHHHHHH |
| BLA-cpRF1-4 | DHPETLVKVKDAEDQLGARVGYIELDLNSGKILESFRPEERFPMMSTFKVLL<br>CGAVLSRVDAGQEQLGRRIHYSQNDLVEYSPVTEKHLTDGMTVRELCSAAI<br>TMSDNTAANLLLTTIGGPKELTAFLHNMGDHVTRLDRWEPELNEAIPNDER<br>DTTMPVAMATTLRKLLTGELLTLASRQQLIDWMEADKVAGPLLRSALPAGW<br>FIADKSGAGERGSRGIIAALGPDGDPDKDGWLDAKEIKALLAAAKAATGAK<br>TIEIEDIPKGGSGGSGGKKVITISEDTPESFEVKMGKGLLKVTVPGKTFIVGKP<br>SRIVVIYTTGSQATMDERNRQIAEIGASLIKHWGKLAAALEHHHHHH |
| BLA-cpRF1-5 | DHPETLVKVKDAEDQLGARVGYIELDLNSGKILESFRPEERFPMMSTFKVLL<br>CGAVLSRVDAGQEQLGRRIHYSQNDLVEYSPVTEKHLTDGMTVRELCSAAI<br>TMSDNTAANLLLTTIGGPKELTAFLHNMGDHVTRLDRWEPELNEAIPNDER<br>DTTMPVAMATTLRKLLTGELLTLASRQQLIDWMEADKVAGPLLRSALPAGW<br>FIADKSGAGERGSRGIIAALGPDGAKTIEIEDIPKGGSGGSGGKKVITISEDT<br>PESFEVKMGKGLLKVTVPGKTFIVEDPDKDGWLDAKEIKALLAAAKAATGK<br>PSRIVVIYTTGSQATMDERNRQIAEIGASLIKHWGKLAAALEHHHHHH |
| BLA-cpRF2-1 | DHPETLVKVKDAEDQLGARVGYIELDLNSGKILESFRPEERFPMMSTFKVLL<br>CGAVLSRVDAGQEQLGRRIHYSQNDLVEYSPVTEKHLTDGMTVRELCSAAI<br>TMSDNTAANLLLTTIGGPKELTAFLHNMGDHVTRLDRWEPELNEAIPNDER<br>DTTMPVAMATTLRKLLTGELLTLASRQQLIDWMEADKVAGPLLRSALPAGW<br>FIADKSGAGERGSRGIIAALGPDGDTLVEISPEEAERLAKTRPVLEVEDPDK |

| | |
|---|---|
| | DGWLDAKERAWILEHLRETHPDASGAVIVVVDGGSGGRRAYLLRVDGKP SRIVVIYTTGSQATMDERNRQIAEIGASLIKHWGKLAAALEHHHHHH |
| BLA-cpRF2-2 | DHPETLVKVKDAEDQLGARVGYIELDLNSGKILESFRPEERFPMMSTFKVLL CGAVLSRVDAGQEQLGRRIHYSQNDLVEYSPVTEKHLTDGMTVRELCSAAI TMSDNTAANLLLTTIGGPKELTAFLHNMGDHVTRLDRWEPELNEAIPNDER DTTMPVAMATTLRKLLTGELLTLASRQQLIDWMEADKVAGPLLRSALPAGW FIADKSGAGERGSRGIIAALGPDGEEAERLAKTRPVLEVEDPDKDGWLDAK ERAWILEHLRETHPDASGAVIVVVDGGSGGRRAYLLRVDPDTLVEISGKPS RIVVIYTTGSQATMDERNRQIAEIGASLIKHWGKLAAALEHHHHHH |
| BLA-cpRF2-3 | DHPETLVKVKDAEDQLGARVGYIELDLNSGKILESFRPEERFPMMSTFKVLL CGAVLSRVDAGQEQLGRRIHYSQNDLVEYSPVTEKHLTDGMTVRELCSAAI TMSDNTAANLLLTTIGGPKELTAFLHNMGDHVTRLDRWEPELNEAIPNDER DTTMPVAMATTLRKLLTGELLTLASRQQLIDWMEADKVAGPLLRSALPAGW FIADKSGAGERGSRGIIAALGPDGVLEVEDPDKDGWLDAKERAWILEHLRE THPDASGAVIVVVDGGSGGRRAYLLRVDPDTLVEISPEEAERLAKTRGKPS RIVVIYTTGSQATMDERNRQIAEIGASLIKHWGKLAAALEHHHHHH |
| BLA-cpRF2-4 | DHPETLVKVKDAEDQLGARVGYIELDLNSGKILESFRPEERFPMMSTFKVLL CGAVLSRVDAGQEQLGRRIHYSQNDLVEYSPVTEKHLTDGMTVRELCSAAI TMSDNTAANLLLTTIGGPKELTAFLHNMGDHVTRLDRWEPELNEAIPNDER DTTMPVAMATTLRKLLTGELLTLASRQQLIDWMEADKVAGPLLRSALPAGW FIADKSGAGERGSRGIIAALGPDGDPDKDGWLDAKERAWILEHLRETHPD ASGAVIVVVDGGSGGRRAYLLRVDPDTLVEISPEEAERLAKTRPVLEVGKPS RIVVIYTTGSQATMDERNRQIAEIGASLIKHWGKLAAALEHHHHHH |
| BLA-cpRF2-5 | DHPETLVKVKDAEDQLGARVGYIELDLNSGKILESFRPEERFPMMSTFKVLL CGAVLSRVDAGQEQLGRRIHYSQNDLVEYSPVTEKHLTDGMTVRELCSAAI TMSDNTAANLLLTTIGGPKELTAFLHNMGDHVTRLDRWEPELNEAIPNDER DTTMPVAMATTLRKLLTGELLTLASRQQLIDWMEADKVAGPLLRSALPAGW FIADKSGAGERGSRGIIAALGPDGDASGAVIVVVDGGSGGRRAYLLRVDPD TLVEISPEEAERLAKTRPVLEVEDPDKDGWLDAKERAWILEHLRETHGKPS RIVVIYTTGSQATMDERNRQIAEIGASLIKHWGKLAAALEHHHHHH |
